# Malaria control and the unexpected spread of diagnostic-resistant *Plasmodium falciparum* in Peru

**DOI:** 10.64898/2026.02.25.707493

**Authors:** Isabela Gerdes Gyuricza, Abebe A. Fola, Alfred Simkin, Kyaw L. Thwai, Jonathan J. Juliano, Jeffrey A. Bailey, Parul Johri, Cobi M. Henry, Luis Cabrera-Sosa, Gerardo Porras-Laymito, Qin Cheng, Oliver J. Watson, Dionicia Gamboa, Hugo O. Valdivia, Jonathan B. Parr

## Abstract

*Plasmodium falciparum* parasites with deletions of the histidine-rich protein 2 and 3 (*hrp2* and *hrp3*) genes evade detection by common rapid diagnostic tests (RDTs) and pose a growing threat to malaria control. While these deletions have emerged in multiple regions globally, the evolutionary forces driving their spread remain unclear. Here, we analyze 1,215 *P. falciparum* samples collected between 2003 and 2018 in Loreto, Peru. This region experienced a major decline in malaria transmission following the Project for Malaria Control in Andean Border Areas (PAMAFRO) and now harbors a high proportion of *hrp2/3* deleted parasites despite limited RDT use. Using molecular inversion probe (MIP) sequencing across > 2,000 genome-wide loci, we observed a marked reduction in genetic diversity, increased clonality, and fixation of parasites with deletions of both *hrp2* and *hrp3* genes (*hrp2-/3-*) over time. Identity-by-descent (IBD) analysis revealed rapid expansion of a single *hrp2-/3-* dominant lineage in the post-PAMAFRO period, consistent with clonal replacement after intense malaria control. Targeted sequencing of the *hrp2/3* regions showed conserved deletion breakpoints across three different lineages, indicative of recombination of a common haplotype into distinct genetic backgrounds. To investigate the evolutionary forces driving the fixation of *hrp2-/3-* in Loreto, we simulated allele frequency trajectories under different selection coefficients. We found that fixation of *hrp2-/3-* due solely to genetic drift (selection coefficient *s* = 0) is unlikely; a selection coefficient of *s* ≥ 0.03 was required for fixation to occur consistently. However, our simulations also indicate that a genetic bottleneck caused by PAMAFRO increased the likelihood of fixation through drift by 4.5- to 17-fold depending on the population. These findings suggest that *hrp2-/3-* fixation was likely driven by a combination of demographic changes resulting from PAMAFRO and selective advantage unrelated to RDT use. Our results demonstrate how intensive malaria control efforts can reshape parasite populations and underscore the value of expanded genomic surveillance as countries move toward malaria elimination.

## INTRODUCTION

In 2024, the World Health Organization (WHO) estimated 610,000 malaria deaths worldwide, the majority of which were caused by the parasite *Plasmodium falciparum*.^1^ In South America, *P. falciparum* transmission has declined substantially over the past two decades in the setting of widespread implementation of malaria control strategies.^2^ However, recent increases in malaria cases in several countries in the Amazon region associated with emerging drug resistance, illegal mining activities, and climate change threaten these gains.^3–5^

In Peru, malaria transmission is largely confined to the Amazon Basin, particularly the Loreto region, where *P. vivax* accounts for more than 80% of cases, though *P. falciparum* transmission continues.^6^ Between October 2005 and December 2010, Peru participated in the Project for Malaria Control in Andean Border Areas (PAMAFRO), a Global Fund-supported initiative that drastically reduced malaria incidence in the region.^7^ However, after the program’s conclusion, malaria cases began to rise again.^7,8^ Previous studies have shown that intensive malaria control efforts in Peru reshaped *P. falciparum* parasite population structure, leading to genetic bottlenecks and clonal replacement.^9,10^ In such settings, reduced transmission allows only a few parasite lineages to survive, which then propagate clonally with minimal recombination. This pattern is evident in the post-PAMAFRO period, marked by a sharp rise in parasites carrying deletions of both histidine-rich protein 2 and 3 (*hrp2* and *hrp3*) genes, a trend likely driven by the clonal expansion of a dominant lineage.^11^

These deletions were first identified in field isolates from Loreto in 2010 and allow parasites to evade detection by HRP2-based rapid diagnostics tests (RDTs).^12^ Parasites with these deletions pose a significant threat to malaria control efforts globally, especially in African countries.^13,14^ Africa accounts for more than 90% of global *P. falciparum* deaths, and most countries rely primarily on RDTs to guide malaria case management.^1^ *Hrp2/3*-deleted parasites could have a selective advantage under widespread RDT use in these countries.^1,13^ In contrast, RDT use is limited and not recommended by WHO for diagnosis of falciparum malaria in Peru. During PAMAFRO, most malaria diagnoses relied on microscopy, with RDTs used mainly in isolated communities.^15,16^ It is therefore unlikely that RDT use drove strong selection for *hrp2/3* deletions in this region. A recent study analyzed *P. falciparum* genomes from samples collected post-2006 in Loreto and found no evidence of positive selection associated with these deletions.^10^ However, the drivers of the increased prevalence of *hrp2/3* deletions remain unclear. Distinguishing between positive selection and recovery from a genetic bottleneck is challenging since both processes can result in similar genomic signatures.^17^ These large deletions, which encompass multiple genes and have been associated with large translocation events, could still confer an unknown fitness advantage.^18^

Understanding how large-scale control interventions like PAMAFRO impact parasite populations is critical for national malaria programs aiming to eliminate malaria and prevent resurgence. While previous studies have focused on the association between control efforts and the emergence of drug resistance, less attention has been paid to the spread of “diagnostic-resistant” strains.^19–21^ Investigating the factors driving the increased prevalence of *hrp2/3* deletions in Peru can help forecast their spread, particularly in settings where similar interventions are implemented and where malaria diagnosis and treatment rely heavily on RDTs, such as Africa. Here, we analyze a 15-year time series of 1,215 *P. falciparum* samples collected in Loreto, Peru, to investigate the population dynamics of *hrp2/3*-deleted parasites. Using *hrp2/3* deletion genotyping and molecular inversion probe (MIP) sequencing, we genetically characterized the parasite population before, during, and after PAMAFRO. We then used simulations to explore potential drivers of the expansion of *hrp2/3*-deleted strains in Peru, highlighting the evolutionary consequences of malaria control on parasite populations.

## RESULTS

### Study population

In this retrospective analysis of samples collected between 2003 and 2018 in previous studies (**Table 1**), we sequenced a total of 1,215 *P. falciparum* samples originating from the Loreto region in the northeastern part of the Peruvian Amazon. Most samples (n = 1,157) were from Maynas province, followed by Loreto (n = 28), Alto Amazonas (n = 16), Requena (n = 14), Datem del Marañón (n = 2), and Mariscal Ramón Castilla (n = 2) provinces (**Figure 1A** and **Supplemental Table 1**). The distribution of samples across years was uneven, with the highest number of samples from 2004 (n = 282), 2005 (n = 168), and 2015 (n = 202) (**Figure 1B** and **Supplemental Table 1**). DNA extracted from dried blood spots (DBS) underwent molecular inversion probe (MIP) sequencing using 2,128 probes targeting genome-wide loci as previously described.^22^ We limited our analysis to 492 samples with high-quality sequencing data (**Extended Data Figure 1**) and 1,692 biallelic SNPs after filtering. These samples originated from Maynas (n = 487) and Alto Amazonas (n = 5). Samples from 2006 and 2007 had inadequate sequencing quality for downstream analysis.

**Figure 1.**
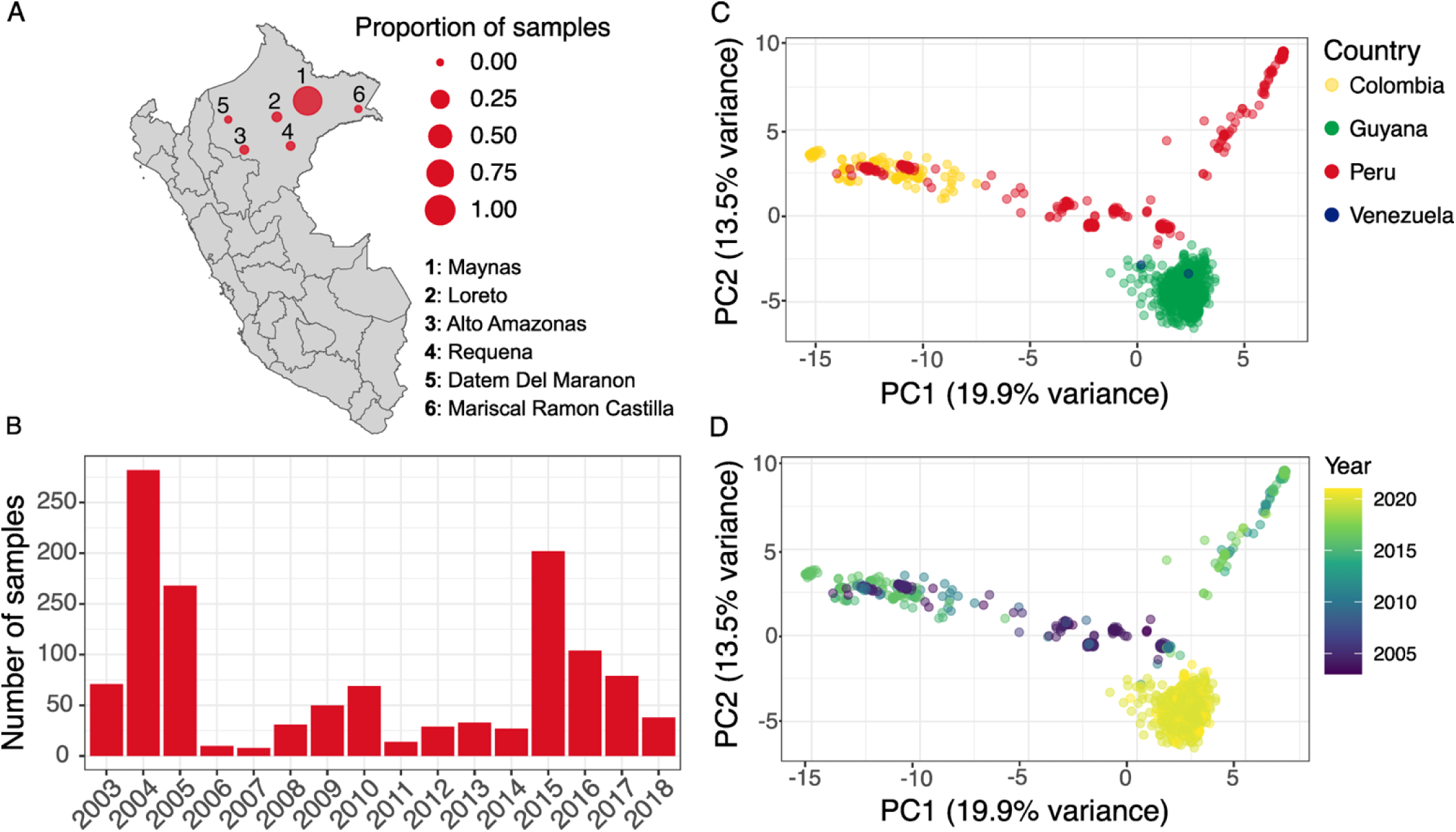
*P. falciparum* samples and population structure in South America. a) Proportion of samples sequenced in this study by province in the Loreto region of northern Peru, originally collected as part of five different studies (n = 1,215). b) Number of samples with high-quality sequencing data included in downstream analysis (n = 492). Principal component analysis (PCA) of allele frequencies from these samples from Peru (n = 492) and publicly available whole-genome sequencing (WGS) data from Colombia (n = 159), Peru (n = 21), Venezuela (n = 2), and Guyana (n = 656) by c) country and d) year of sample collection.

**Table 1.** Description of sample sets included in this study.

| Study label | Reference | Province | Year of collection | Study population |
| --- | --- | --- | --- | --- |
| Artekin | Grande et al., 2007<sup>44</sup> | Maynas | 2003-2005 | Includes individuals aged 5–60 years presenting with fever and confirmed *P. falciparum* infection. |
| PAMAFRO | Bendezu et al., 2010<sup>45</sup> | Maynas | 2006 | Passive malaria case detection includes individuals with clinical symptoms suspicious of malaria. |
| FIND | Gamboa et al., 2010<sup>12</sup><br>Bendezu et al., 2022<sup>46</sup> | Maynas, Alto Amazonas, Loreto, Requena and Mariscal Ramón Castilla | 2007-2010, 2014, and 2017 | Active and passive malaria case detection includes individuals older than 5 years with confirmed *P. falciparum* infection. |
| Círculo estudio corto | Moreno-Gutierrez et al., 2018<sup>47</sup> | Maynas | 2015 | Population-based cohort study. |
| NAMRU | Valdivia et al., 2022<sup>11</sup> | Maynas | 2011-2018 | Passive surveillance protocol includes individuals older than 1 year old, with fever, and suspected or confirmed malaria. |

### *P. falciparum* population structure in South America

Given the limited literature on *P. falciparum* population structure in South America, we first explored how Peruvian parasite populations cluster with isolates from other South American countries. Publicly available whole-genome sequencing (WGS) data from MalariaGEN Pf7^23^ (Colombia, n = 159, years 2011-2017; Peru, n = 21, years 2009 and 2011; Venezuela, n = 2, years 2012 and 2016) and Schwabl et al.^24^ (Guyana, n = 656, years 2020 and 2021) were downloaded and merged with the Peru MIP dataset. Analysis was limited to 1,366 high-quality SNPs that were shared among all data sets.

Principal component analysis (PCA) revealed clustering by geography (**Figure 1C**). Within Peru, parasites segregated into two distinct temporal groups (**Figure 1D**). Peruvian parasites collected before 2011 showed greater genetic similarity to Colombian (collected between 2011 and 2017) and Guyanian (collected in 2020 and 2021) parasites. Peruvian samples collected after 2011, when the PAMAFRO malaria control initiative concluded, formed a distinct cluster.

### Clonal expansion of *hrp2-/hrp3-* parasites in Peru

*Hrp2/3* deletion genotyping by multiplex real-time PCR confirmed rising frequency of *hrp2*-/*hrp3*- (*hrp2*-/*3*-) parasites in Peru over time. A high proportion of parasites carried only *hrp3* deletions (*hrp2*+/*hrp3*-) until 2008, peaking at 46.5% - 60% between 2003 and 2006 (**Figure 2A**), with a shift toward *hrp2-/3-* parasites occurring after 2008-2010. The proportion of *hrp2-/3-* parasites rose from 18.3% (13 of 71 samples) in 2003 to 86.8% (33 of 38 samples) by 2018. Additional genotyping results are described in the supplemental results and **Supplemental Tables 2-4**.

**Figure 2.**
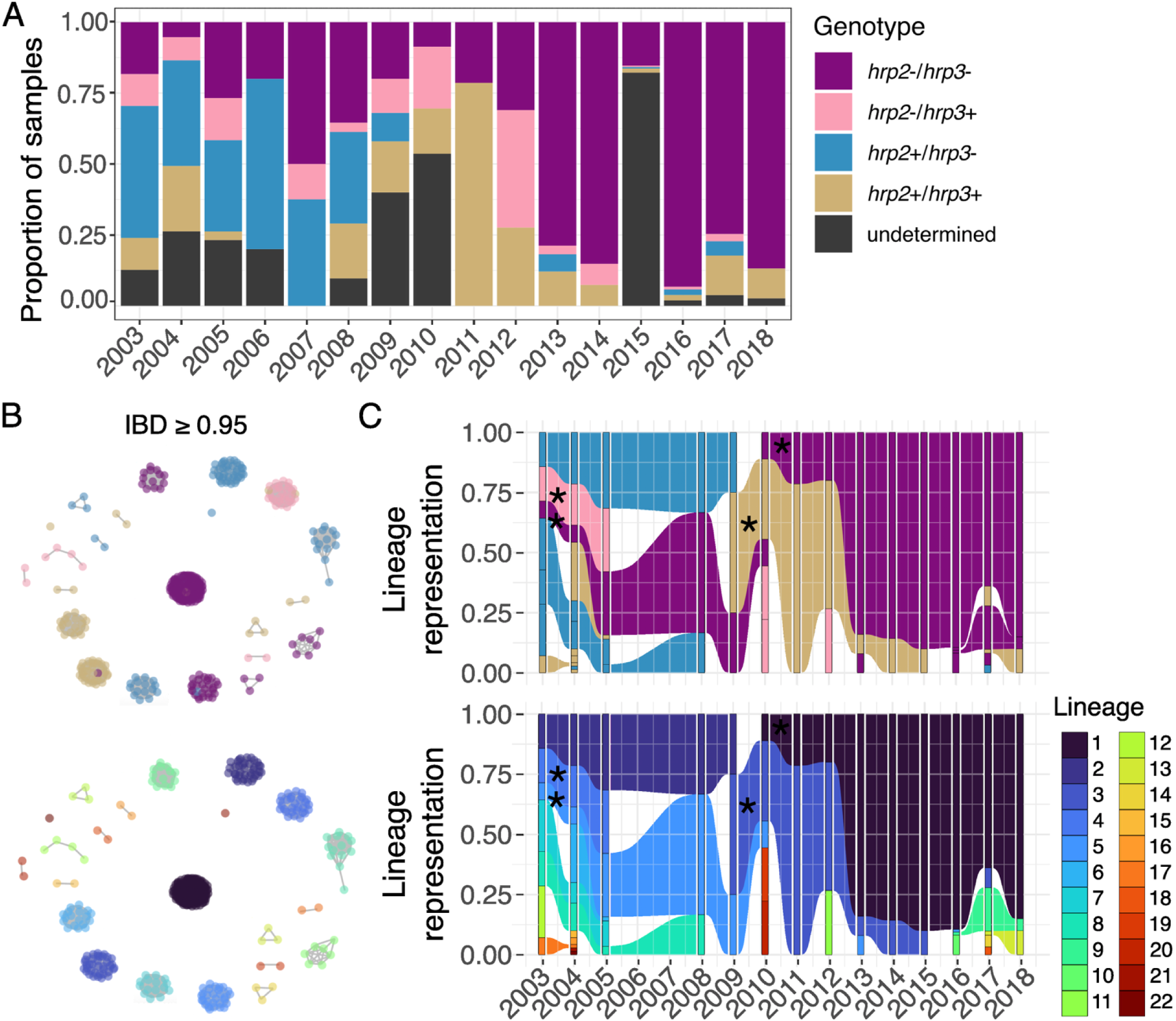
Clonal expansion of *hrp2*-/*3*- parasite populations in Loreto, Peru. a) Proportion of samples with each *hrp2/3* genotype per year as estimated by PCR (n = 1,215). Samples of undetermined genotype failed to amplify the human beta-tubulin gene and/or had insufficient parasite DNA as determined by multiplexed PCR. b) Identity-by-descent (IBD) network analysis showing 22 genetically distinct, clonal (IBD ≥ 0.95) lineages of parasites (n = 492). Each dot represents a parasite, colored by *hrp2/3* genotype (top) and lineage (bottom). c) IBD lineage representation over time, colored by *hrp2/3* genotype (top) and lineage number (bottom). *Lineages containing parasites with more than one *hrp2/3* genotype. Rectangles represent distinct lineages by year; shaded areas illustrate lineages that persisted over time.

MIP sequencing confirmed clonal expansion of *hrp2-/3-* strains after PAMAFRO. Using high-coverage genome-wide MIP sequencing data generated for 492 samples, we assessed genetic relatedness among parasites through identity-by-descent (IBD) analysis. First, we estimated the complexity of infection (COI), and found that all the genotyped samples were monogenomic (COI = 1). We then filtered the genotype data to include only samples with *hrp2/3* deletion calls, resulting in a dataset of 467 samples. Using an IBD threshold of ≥ 0.95 to define highly related (i.e., clonal) parasites, our analysis identified 22 genetically distinct clusters of clonal parasites circulating in Peru between 2003 and 2018. These clusters are described throughout as lineages and comprise 92.3% (431 of 467) of parasites analyzed.

Parasite lineages largely clustered by *hrp2/hrp3* deletion status (**Figure 2B, C - top**). Although four lineages displayed mixed genotypes, more than 90% of parasites within each lineage consistently maintained the same *hrp2/3* profile over time (**Supplemental Table 5**). Among parasites with the *hrp2+/hrp3-* genotype (i.e., *hrp3*-), four distinct lineages were observed in 2003. Two of these, lineages 2 and 8, persisted until 2009 and 2008, respectively, while two *hrp3*- samples from a separate lineage (lineage 17) appeared later, in 2017 (**Figure 2B, C - bottom**). Parasites with the *hrp2-/hrp3+* genotype (i.e., *hrp2*-) detected between 2003 and 2005 all belonged to lineage 4. While these may have persisted beyond 2005, they were not included in the IBD analysis due to insufficient quality sequences from 2006-2007 (**Extended Data Figure 2**). Alternatively, the *hrp2*- observed after 2005 may have independent origins, supported by their occurrence in distinct lineages in 2010 (lineages 19 and 20) and 2012 (lineage 11).

All 24 *hrp2-/3-* parasites successfully sequenced between 2003 and 2009 belonged to lineage 5, though most samples from 2003 lacked sufficient sequencing coverage (**Extended Data Figure 2**). A notable shift occurred in 2010, when a *hrp2-/3-* sample from a different lineage (lineage 1) was identified. In subsequent years, five additional *hrp2-/3-* lineages (10, 5, 9, 14, and 13) emerged, but lineage 1 rapidly expanded and accounted for 88.2% (201 of 228) of *hrp2-/3-* samples collected after 2010 (**Figure 2C**).

Using a MIP panel targeting *hrp2*, *hrp3*, and their flanking regions, we found that *hrp2-/3-* parasites from Peru shared similar deletion breakpoints (**Extended Data Figure 3 and 4**) that are distinct from those recently observed in Ethiopia.^13^ These data were available for a subset of 123 samples, all collected between 2013 and 2018. We observed strong concordance between *hrp2/3* deletion calls by MIP sequencing and real-time PCR (**Extended Data Figure 3 and 4**). Only two samples originally genotyped as *hrp2+/hrp3*- by PCR exhibited no sequencing coverage across both *hrp2* and *hrp3* loci. Notably, IBD analysis identified these two samples as lineage 1, which is otherwise composed exclusively of *hrp2-/3-* parasites (**Supplemental Table 5** and **Extended Data Figure 3 and 4**), suggesting that these samples were likely misclassified by PCR. Read mapping revealed a consistent chromosomal breakpoint pattern across all Peru samples, including those from distinct IBD lineages (**Extended Data Figures 3 and 4**), suggesting that the *hrp2-/3-* haplotype likely originated in lineage 1 and was later introduced into other lineages through recombination. Deletions were approximately 35.3 kb (1,367,511 - 1,402,810) for chromosome 8 and 44.8 kb (2,808,674 - 2,853,444) for chromosome 13, encompassing multiple genes. To assess the specificity of the breakpoint patterns, we also compared the Peruvian deletion profiles with those of parasites from Ethiopia, previously generated using the same *hrp2/3* MIP panel.^13^ The analysis revealed distinct breakpoints on chromosome 8 (*hrp2*) in each country (**Extended Data Figure 5**). We did not find differences in breakpoints between Peru and Ethiopia on chromosome 13 (*hrp3*), but MIP target density was limited (**Extended Data Figure 6**).

### Shifts in parasite genetic structure associated with PAMAFRO malaria control

To investigate temporal changes in population structure among *P. falciparum* isolates from Peru, we performed admixture analysis on high-quality genotype data obtained from MIP sequencing of 492 samples. To explore how parasite genetic structure changed over time in relation to the PAMAFRO malaria control program, we grouped samples into three time periods: pre-PAMAFRO (before 2006), during PAMAFRO (2006–2011), and post-PAMAFRO (after 2011). To assess how genetic structure varied by *hrp2/3* genotype, samples were also stratified according to their deletion status.

Across all *hrp2/3* genotypes, parasites exhibited shifts in genetic profiles during or after the PAMAFRO period (**Figure 3**). Parasites carrying deletions of only *hrp2* or *hrp*3 were more common prior to 2006, and showed higher levels of admixture and genetic diversity. This is consistent with multiple co-circulating lineages identified in the IBD analysis (**Figure 2B and C**). Although these parasite populations underwent changes in composition post-PAMAFRO, they continued to display signs of admixture, suggestive of recombination between genetically distinct strains carrying single deletions. In contrast, *hrp2-/3-* parasites exhibited markedly lower admixture over time (**Figure 3**). A shift in the population structure of *hrp2-/3-* parasites occurred between the pre- and post-PAMAFRO periods, with *hrp2-/3-* parasites from each period characterized by a single dominant ancestry component. These findings are consistent with IBD results, highlighting the replacement and clonal expansion of a *hrp2-/3-* lineage following the decline of malaria transmission in Peru.

**Figure 3.**
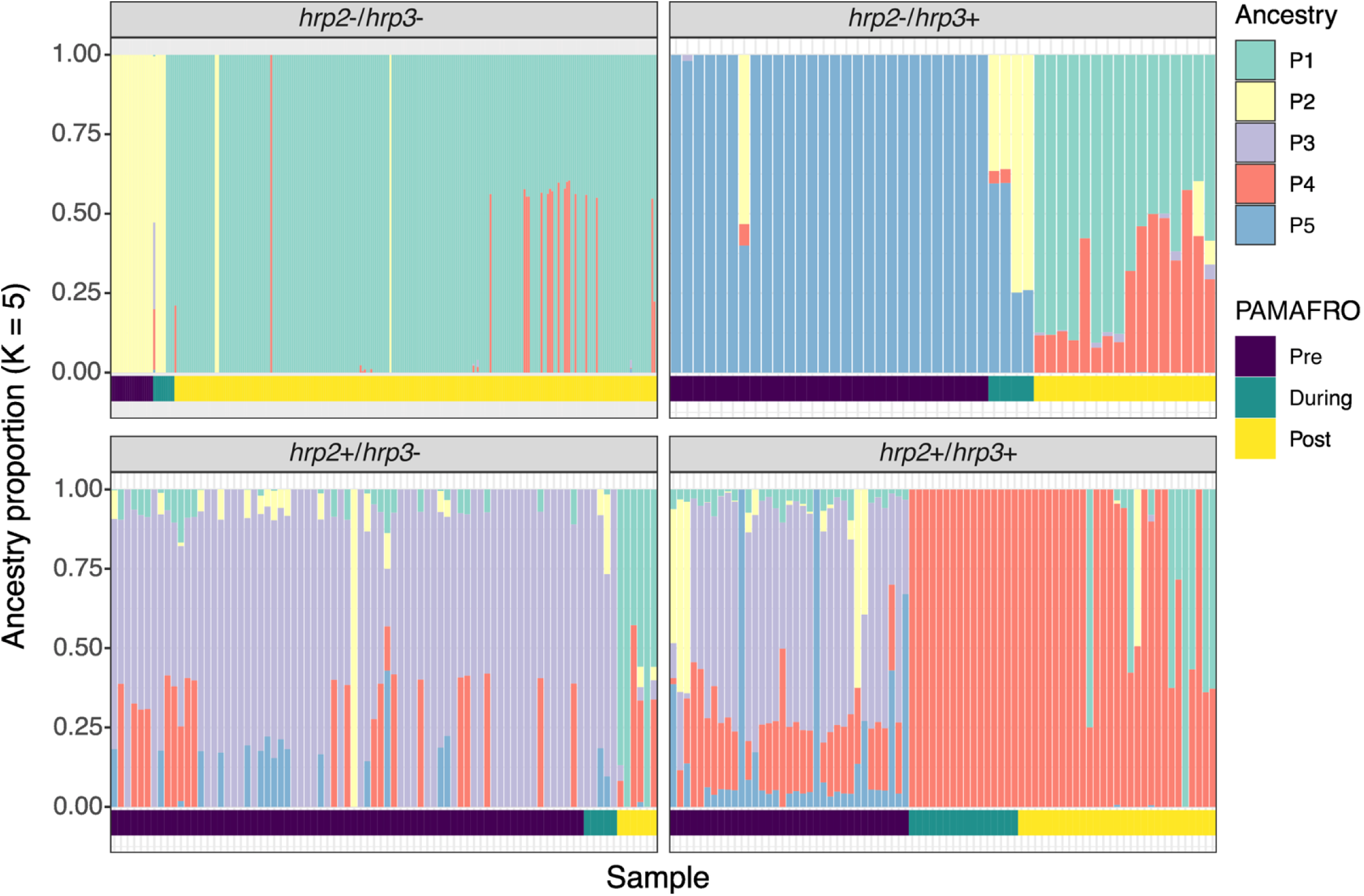
Genetic structure of parasites before, during, and after PAMAFRO. Ancestry proportions inferred using sparse non-negative matrix factorization (sNMF) for the optimal number of K = 5 ancestral populations (n = 492). Each bar represents an individual parasite, with colors indicating ancestry components. Parasites are grouped by *hrp2/3* genotype and sorted by PAMAFRO period: pre-PAMAFRO (samples collected before 2006), during-PAMAFRO (2006–2011), and post-PAMAFRO (after 2011).

### Genome-wide reduction in nucleotide diversity after PAMAFRO malaria control

Our results show changes in admixture and parasite structure associated with PAMAFRO, and its impact on reducing malaria prevalence is well documented.^7^ We sought to quantify this effect by examining changes in effective population size over time. We computed genome-wide nucleotide diversity (π) using samples with high-coverage MIP sequencing data, focusing on synonymous SNPs and invariant regions. Because genetic diversity and effective population size are directly related, changes in π serve as a proxy for fluctuations in effective population size.

Our analysis revealed a marked decrease in nucleotide diversity and, consequently, in effective population size following the PAMAFRO intervention period (**Extended Data Figure 7A**). To account for variation in sample size over time, we repeated the analysis using a rarefied random subset of 18 samples per year, the minimum available across timepoints. The rarefied results confirmed the observed decline in diversity after 2011, indicating that the trend is not an artifact of uneven sampling (**Extended Data Figure 7B**). We next evaluated whether the reduction in diversity was driven by specific genomic regions by computing π separately for each chromosome. Our results show a genome-wide reduction in diversity following PAMAFRO (**Figure 4A**). Changes in minor allele frequency (MAF) distributions over time provide additional evidence of loss of diversity. Prior to 2011, MAF distributions were broader and skewed toward intermediate frequencies, reflecting a genetically diverse parasite population (**Figure 4B**). After 2011, most SNPs shifted toward very low frequencies, with peaks near fixation (**Figure 4B**). These patterns are consistent with loss of genetic diversity after a bottleneck followed by selection after PAMAFRO. To investigate signatures of recent positive selection in our data, we attempted integrated haplotype score (iHS) analyses. However, reliable iHS estimates across the genome were not possible owing to the low and uneven density of informative SNPs in the population.

**Figure 4.**
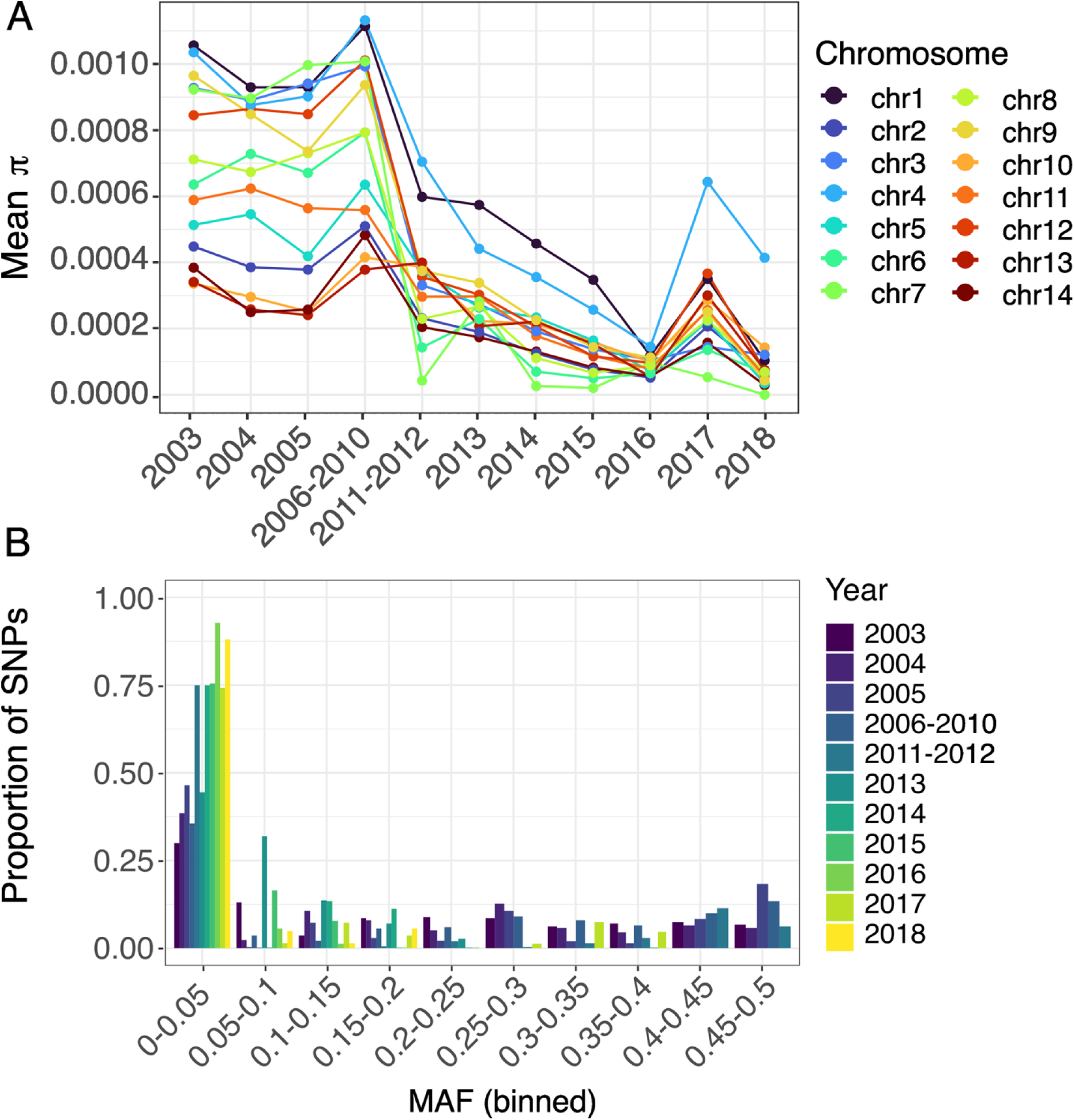
Genome-wide changes in nucleotide diversity and minor-allele frequency following PAMAFRO. a) Mean genome-wide nucleotide diversity (π) per sample (n = 492), calculated across non-overlapping 5 kb windows using synonymous and invariant sites. Mean π per chromosome and per year group was computed by summing the observed nucleotide differences across all windows and dividing by the total number of comparable (callable) sites. Samples are grouped by year of collection; years with fewer than 18 samples were combined with adjacent years to ensure adequate representation. b) Minor allele frequency (MAF) distributions for each year group. Allele frequencies were binned into 20 equal-width intervals (0–0.5) to visualize genome-wide trends.

### The impact of PAMAFRO and population size on the fixation of *hrp2/3* deletions

To assess whether the fixation of *hrp2/3* deletions after PAMAFRO could be explained by genetic drift following a bottleneck, we used two complementary simulation approaches.

First, we used SLiM^25^ to directly model *hrp2* and *hrp3* allele frequency trajectories over a 15-year period in the *P. falciparum* population, informed by annual falciparum malaria incidence data reported by CDC Peru.^8^ We modeled 1000 independent replicate simulations over time (assuming 9 parasite generations per year) and using three different population size scenarios with parasite population size estimated based on: (i) all reported cases nationwide; (ii) all reported cases from the province of Maynas; and (iii) reported cases from the districts most represented in our dataset (Iquitos, Punchana, and San Juan Bautista; **Supplemental Table 1**) (**Figure 5A**). We found that when no positive selection was assumed for deletions of *hrp2* or *hrp3* (selection coefficient, *s* = 0), only a small proportion of simulations (0-10%) reached *hrp2-/3-*fixation (defined as >75% frequency) by 2018, with the highest proportion (9.3%) observed at the district level (**Figure 5B**). To observe fixation of *hrp2-/3-* in most simulated scenarios, a selection coefficient of at least 0.03 was required (**Figure 5B**). Additionally, the observed frequency of *hrp2-/3-* diverges most from the simulated neutral trajectories after 2012, suggesting that positive selection likely began around this time, coinciding with the completion of the PAMAFRO program (**Extended Data Figure 8**). Even under moderate selection (*s* ≥ 0.03, corresponding to *Ns* ≥ 15, where *N* is the haploid effective population size), some simulations at the district and province levels produced intermediate allele frequencies of *hrp2-/3-* rather than fixation (**Figure 5B**). This underscores the impact of stochastic processes such as genetic drift in smaller populations. These findings demonstrate the influence of population size on the frequency of *hrp2/3* deletions in Peru and suggest that moderate selective advantage is necessary for *hrp2-/3-* to reach fixation in population sizes reflective of the province of Maynas or larger.

**Figure 5.**
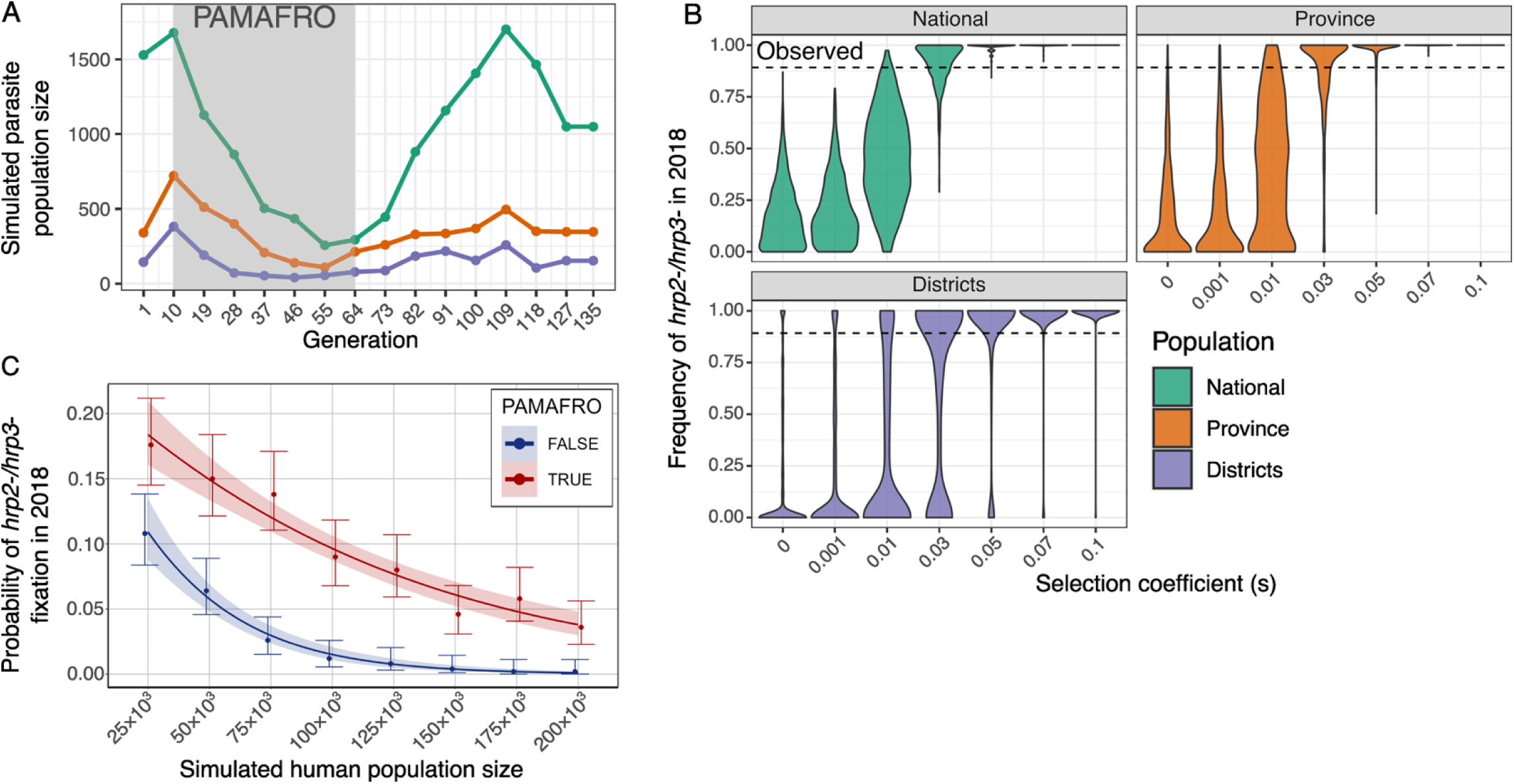
Simulated probability of *hrp2-/3-* fixation following PAMAFRO. a) Number of parasites simulated per generation in SLiM, based on annual *P. falciparum* case reports from CDC Peru for three spatial scales: (i) all reported cases nationwide (National); (ii) reported cases for the province of Maynas (Province); and (iii) reported cases from Iquitos, Punchana, and San Juan Bautista (Districts). b) Simulated frequency of *hrp2-/3-* in 2018 under varying selection coefficients (s), using SLiM. The dashed line represents the observed frequency of *hrp2-/3-* in our dataset from 2018. c) Probability of *hrp2-/3-* fixation (frequency ≥ 0.75) across different human population sizes, simulated using a *P. falciparum*-specific malaria transmission model. Simulations were run both with (red) and without (blue) the PAMAFRO-driven population bottleneck.

Next, we used a *P. falciparum*-specific mathematical model of malaria transmission and parasite genetics.^26^ This implementation explicitly tracks individual humans, mosquitoes, and parasite clones, enabling realistic parasite population bottlenecks during transmission and explicit modelling of how transmission and intervention dynamics shape parasite genetic variation. We calibrated the model to the per-capita *falciparum* malaria case numbers reflective of the same three districts (Iquitos, Punchana, and San Juan Bautista) and ran 500 simulations across varying human population sizes (25,000 - 200,000), comparing scenarios with and without the PAMAFRO-driven reduction in malaria incidence. As expected from population genetic theory and observed in our SLiM modeling exercise, we observed higher frequencies of *hrp2-/3-* due to drift in smaller population sizes (**Extended Data Figure 9**). To quantify the impact of PAMAFRO across population sizes, we fit a binomial regression model to the simulation outputs, modelling the probability of *hrp2-/3-* fixation as a function of population size and PAMAFRO. We find that PAMAFRO significantly increased the odds of *hrp2-/3-* fixation by modifying the strength of genetic drift across population sizes (p = 4.07 x 10^-9^; **Figure 5C**). In human populations of N = 150,000, comparable to the population size of the Iquitos district prior to PAMAFRO, PAMAFRO increased the odds of *hrp2-/3-* fixation by 17-fold (95% CI: 9.3–31.0), corresponding to fixation in approximately 6% of simulations (**Figure 5C**). In smaller populations of approximately N = 75,000, comparable to San Juan Bautista, PAMAFRO increased the odds of fixation by 4.5-fold (95% CI: 3.5–5.8), with fixation occurring in more than 12% of simulations. Overall, without PAMAFRO, the probability of *hrp2-/3-* fixation did not exceed 5% except in very small populations (N = 25,000), whereas under PAMAFRO this threshold was exceeded even at realistic population sizes of approximately 150,000.

Taken together, these complementary modeling approaches suggest that *hrp2-/3-*parasites survived a genetic bottleneck caused by PAMAFRO and expanded thereafter in the setting of a selective advantage conferred by the deletions.

## DISCUSSION

Our study uncovers profound shifts in the genetic landscape of *Plasmodium falciparum* in Peru, with the most dramatic changes emerging after 2010–2012 and coinciding with the cessation of the PAMAFRO malaria control initiative. Leveraging a robust longitudinal dataset, we provide the most comprehensive genomic snapshot to date of this transition. We reveal a collapse in genetic diversity, a surge in clonal lineages, and fixation of parasites harboring deletions of both *hrp2* and *hrp3*. Using simulations, we show that this fixation is unlikely to have resulted from genetic drift alone, although the sharp population decline following PAMAFRO increased the likelihood of drift-driven outcomes. Instead, a selective advantage probably also contributed to the rise of *hrp2-/3-* parasites in the region. These results underscore how large-scale control efforts can reshape parasite populations and inadvertently drive the rise of diagnostic-resistant lineages.

Before 2012, the *P. falciparum* population in Loreto exhibited substantial genetic diversity, higher minor allele frequencies, and multiple circulating lineages, including parasites with single *hrp2* or *hrp3* deletions, corroborating previous reports.^10,27,28^ Starting in 2013, the parasite population became increasingly dominated by a single *hrp2-/3-* lineage (lineage 1), which persisted and expanded in subsequent years. This lineage likely corresponds to lineage “155” described by Kattenberg et al.^27^, lineage “Bv1” from Villela et al.^28^, and haplotype “H8” reported by Valdivia et al.^11^ These findings align with a recent study showing the reduction in the number of *P. falciparum* populations in Loreto after PAMAFRO, along with a decline in genetic diversity likely driven by limited recombination in a low-transmission setting.^10^ Although we could not generate *hrp2/3*-targeted MIP data from samples before 2013, our data show that *hrp2-/3-* samples from three different lineages (lineages 1, 9, and 10) carry identical deletion breakpoints, suggesting that the same *hrp2/3*-deleted haplotype was inherited from a common ancestral lineage and spread through recombination into a few genetically distinct backgrounds.

Deletions of *hrp2* and *hrp3* have now been reported across multiple countries, mostly in South America and the Horn of Africa.^29^ Although the precise mechanisms behind the origin of these deletions are not fully resolved, studies have shown that they are common in both field and laboratory strains and have arisen independently across different genetic backgrounds.^18,30^ These deletions are thought to result from genomic processes such as meiotic misalignment, non-allelic homologous recombination, and telomere healing.^18^ *In vitro* studies have shown that deletions of both *hrp2* and *hrp3* can reduce parasite fitness.^31^ However, real-world epidemiological dynamics may overcome these disadvantages under certain conditions. For example, in Ethiopia, we previously reported that *hrp2* deletion appears to be under strong positive selection, likely driven by widespread RDT use to guide treatment decisions.^13^ In Peru, selection of *hrp2-/3-* by RDT-driven treatment decisions is unlikely. During PAMAFRO, most diagnostic effort was focused on microscopy, but RDTs were also distributed.^15^ The province of Maynas reported using 26,855 RDTs (including both HRP2 and LDH-based) for malaria testing, with the highest use occurring in 2008.^15^ These were implemented mostly in isolated communities located at least one hour from the closest microscopy point.^15^ Since 2010, after reports of high numbers of *hrp2-/3-* parasites in Loreto, HRP2-based RDTs were discontinued.^12,32^ This diagnostic history does not align with the timing of positive selection inferred from our simulations, which suggests that selection began around 2012.

Our simulations suggest that a selection coefficient of at least 0.03 is required for these deletions to approach fixation, as observed. We and others^10^ did not find evidence of positive selection surrounding *hrp2/3* deletions, though this moderate strength of selection (with N*s* ≥ 15) is unlikely to generate obvious selective sweeps on genomic scans.^33^ The biological functions of *hrp2* and *hrp3* remain unclear, and it is therefore possible that their deletion confers a selective advantage unrelated to RDT pressure. Another point to consider is that these deletions are large, and a fitness advantage may also result from the concurrent loss of other linked genes. For example, these deletions encompass *stevor* and genes from the *Plasmodium* helical interspersed subtelomeric (PHIST) family, which have been implicated in red blood cell invasion.^13,18,38,39^ In Peru, *hrp3* deletions have also been associated with duplication and translocation of large segments of chromosome 11, which may confer additional benefits to parasite survival.^18^

It is possible that fixation of *hrp2*-/3- parasites was driven by positive selection acting on other loci, such as those associated with antimalarial drug resistance. During PAMAFRO, artemisinin-based combination therapies (ACTs) became widely used in the region.^40^ However, no validated mutations in the propeller region of *kelch13* associated with artemisinin partial resistance have been reported in Peru to date.^27,41^ The sparsity and heterogeneity of SNP spacing in our MIP dataset limited our ability to detect signatures of selection across the genome. Further investigation, including *in vitro* competition assays and clinical studies are required to understand how these complex structural variants affect parasite fitness.

In addition to the role of positive selection, our simulations showed that the genetic bottleneck caused by PAMAFRO increased the likelihood of *hrp2-/3-* fixation, particularly in small populations. Previous studies have shown that fixation of *hrp2-/3-* in Loreto is geographically limited^42^, supporting the idea that stochastic events such as genetic drift could have occurred in isolated pockets with small *P. falciparum* populations. In such settings, a population bottleneck could rapidly reduce genetic diversity and, consequently, competition among parasite strains. Coupled with reduced host immunity, selective advantage, and other unknown factors, this scenario could allow *hrp2-/3-* lineages to persist and expand. Taken together, our modeling results are most consistent with a scenario in which random parasites survived through a population bottleneck, and a moderate selective advantage conferred by the deletions contributed to their spread thereafter.

This study is not without limitations. First, we analyzed samples collected from multiple studies conducted for other purposes. While this reduces generalizability, the breadth of timepoints included in our analysis provides unique opportunities to investigate recent, dramatic shifts in Loreto’s parasite population. Second, models cannot completely recapitulate the many forces impacting parasite evolution, despite our efforts to ground simulations on empirical data. Due to limited information on the effective population size and population structure of *P. falciparum* in Peru, we assumed that reported malaria incidence across districts serves as a reasonable proxy for parasite population size in SLiM. The malaria transmission modelling approach was calibrated using this data with realistic human population sizes for the districts. Both simulation approaches assume that malaria prevalence and the impact of PAMAFRO is homogenous across the district’s population. These assumptions probably oversimplify real-world geographic heterogeneity. Nonetheless, efforts to understand the demographic history of *P. falciparum* populations are essential for interpreting the evolutionary forces shaping parasite genomes.

Our findings and complementary modeling approaches show that drivers of *hrp2-/3-*fixation in Loreto are complex, involving both reduced genetic diversity resulting from the PAMAFRO associated population bottleneck and a selective advantage linked to these deletions. We also show the potential of region-specific, longitudinal genomic data to anticipate and mitigate the unintended consequences of malaria control efforts. As countries approach malaria elimination, parasite populations become smaller and increasingly susceptible to stochastic changes. In this context, genomic surveillance can enable early detection of dangerous lineages while improving understanding of the evolutionary dynamics shaping parasite populations.

## METHODS

### Sampling and study sites

Dried blood spots (DBS) were obtained from 1,215 individuals with confirmed *P. falciparum* infection by microscopy or PCR. Samples were collected through both active and passive malaria case detection across five distinct studies conducted in the Loreto region of Peru between 2003 and 2018, as shown in Table 1. Details of these parent studies are reported elsewhere; all received research ethical committee approval and obtained informed consent from participants. DNA was extracted from three 6 mm punches per DBS using the Chelex-Tween 20 method, as previously described.^43^

### *Hrp2/3* deletion status

Deletions in the *hrp2* and *hrp3* genes were assessed using a multiplexed real-time PCR assay, following methods previously described by Grignard et al. (2020)^48^. Briefly, extracted DNA was amplified using specific primers and probe sets targeting exon 2 of *hrp2* and *hrp3*. Control targets included the *P. falciparum* lactate dehydrogenase (*pfldh*) gene and the human beta-tubulin gene. Human DNA and the laboratory strains 3D7 (*hrp2+/hrp3+*), DD2 (*hrp2-/hrp3+*), and HB3 (*hrp2+/hrp3-*) served as controls. Cycle threshold (Ct) values obtained from real-time PCR assays were analyzed and deletion calls made using custom R scripts. In summary, samples were deemed suitable for deletion analysis if both human beta-tubulin and *pfldh* amplified within a Ct range of 15 to 35. Deletion status was determined by comparing Ct values for *hrp2*/*hrp3* relative to control genes. Samples showing no amplification of *hrp2*/*hrp3* but successful amplification of control genes were classified as deletions, while those showing amplification for both target and control genes were classified as *hrp2*+/*3*+ (wild-type). Samples were manually reviewed and called if *hrp2*/*hrp3* amplification yielded Ct values below 15, above 35, or if the Ct values for *hrp2* or *hrp3* were more than three cycles higher than those obtained for *pfldh*.

### MIP sequencing data

For population genetic analysis, samples were sequenced using a genome-wide MIP panel targeting 2,128 probes distributed across diverse regions of the *P. falciparum* genome, covering at least 50% of the SNPs with minor allele frequency (MAF) > 0.05.^22^ To visualize chromosomal breakpoints and confirm *hrp2/3* deletions, a second MIP panel consisting of 241 probes targeting *hrp2* (chromosome 8), *hrp3* (chromosome 13), and their flanking regions was also used.^13^ MIP capture and library preparation were performed as previously described^22^ and sequencing was done in Illumina NextSeq 550 (150 bp paired-end reads) at Brown University (Rhode Island, USA). To assess assay sensitivity across parasite loads, we also included four control aliquots of the 3D7 laboratory strain at defined parasite densities (500, 1,000, 2,000, and 4,000 parasites/µL). We were only able to generate high-quality *hrp2/3* MIP data for 123 samples collected between 2013-2018. The samples spanned a wide range of parasite densities. No clear correlation between parasite density and MIP coverage was observed, suggesting that sample age was the primary factor influencing *hrp2/3* MIP performance. The sequencing data were processed using the *miptools* pipeline (v0.5.0) as instructed in https://miptools.readthedocs.io/en/latest/. This software uses unique molecular identifiers (UMIs) to extract reads and remove PCR duplicates^49^. Reads were clustered into haplotypes separately for each sample and each MIP probe using a haplotype clustering algorithm. Haplotypes were aligned to the *P. falciparum* 3D7 reference genome (PlasmoDB-42_Pfalciparum3D7) using *BWA*^50^ (v0.7.17). Variant calling was performed using *FreeBayes*^51^ (v1.3.6) with optimized settings for *P. falciparum* including: pooled continuous mode (‘--pooled-continuous’), minimum alternate allele fraction of 0.01 (‘--min-alternate-fraction 0.01’), minimum alternate allele count of 2 (‘--min-alternate-count 2’), haplotype length of 3 (‘--haplotype-length 3’), minimum alternate total of 10 (‘--min-alternate-total 10’), and utilization of the best 70 alleles (‘--use-best-n-alleles 70’). VCF files underwent additional filtering using *bcftools*^52^ (v1.9). Variants were filtered to include only high-quality (quality score > 20) biallelic SNPs. Samples with over 50% missing genotype data were excluded from further analysis. Additionally, samples were filtered based on a minimum depth of coverage (≥ 5X), which was defined by the number of UMI counts divided by the total number of probes in the panel. For the *hrp2/3*-targeted MIP panel, coverage was calculated using UMI counts from probes located upstream of the known deletion breakpoints —specifically, those mapping before position 1,367,332 on chromosome 8 and position 2,807,022 on chromosome 13 to avoid bias introduced by missing data within the deleted regions. Before downstream analysis, samples had their complexity of infection (COI) estimated by the R package COIAF^53^, and no mixed infections (COI > 1) were found.

### Publicly available WGS data

Publicly available WGS data of *P. falciparum* isolates from South America were downloaded from Pf7^23^ and Schwabl et al^24^. Alignment and variant calling were performed using a *GATK4* pipeline optimized for *P. falciparum* as previously described^54,55^. The resulting VCF file underwent the same filtering protocol applied to the MIP data. Additionally, variants were further filtered to retain only genomic positions included in the genome-wide MIP panel from our Peruvian data, facilitating subsequent merging of the two datasets for downstream analysis. The VCF file was further processed in R using the package vcfR^56^ (v1.15.0), and within-sample allele frequencies (WSAF) were estimated by dividing the reference allele read count by the total depth of coverage for each SNP. To address missing data in the WSAF matrix, loci were imputed using the median allele frequency across all samples for each SNP, following the approach implemented in the ‘get_wsaf()’ function from the *MIPanalyzer*^57^ R package (v1.1.0). For comparison with the MIP data, the VCF file was filtered to retain only genomic positions present in the MIP VCF using the ‘--targets-filè option in *bcftools view.* The filtered VCF was then merged with the MIP VCF using *bcftools merge*. Principal component analysis (PCA) was conducted using the ‘pca_wsaf()’ function from the *MIPanalyzer*^57^ package to highlight population structure among isolates from South America.

### Genetic relatedness

Genetic relatedness among *P. falciparum* isolates was assessed using identity-by-descent (IBD) analysis. IBD was calculated based on genotype data extracted from the VCF files using the R package vcfR.^56^ Pairwise IBD values were computed using the function ‘inbreeding_mle()’ from the R package *MIPanalyzer.*^57^ This function implements a maximum-likelihood approach as described by Verity et al. (2020).^58^ Isolates were defined as clonal pairs if their pairwise IBD values were ≥0.95. The resulting genetic relatedness networks were visualized using the R package *ggraph*^59^ (v2.2.1) employing the Fruchterman-Reingold (‘fr’) clustering algorithm.

### Population structure

Population structure was inferred using sparse non-negative matrix factorization (sNMF) as implemented in the R package *LEA*^60^ (v3.9.9). The sNMF algorithm was run for values of K (number of ancestral populations) ranging from 1 to 10, with 5 replicates per K to assess model stability. The optimal number of K was determined based on the minimum cross-entropy criterion across replicates. The lowest cross-entropy was observed at K = 5, which was selected as the best-fitting model to describe the population structure. Results were used to interpret patterns of population structure and admixture over time and *hrp2/3* deletion status.

### Genetic diversity

To estimate nucleotide diversity, VCF files were generated using FreeBayes^51^ within the MIPtools framework, including the same parameters described previously, with the addition of the ‘--report-monomorphic’ flag to retain invariant genomic positions. To account for temporal variation in sample availability, a minimum threshold of 18 samples per year was established, corresponding to the number of samples available in 2003. When fewer than 18 samples were available in a given year, adjacent years were combined to meet this threshold. The resulting VCF was split into two datasets: one containing only monomorphic sites, which was filtered using *bcftools*^50^ to retain only sites with read depth ≥10 and genotype data in at least 50% of samples; and a second containing biallelic SNPs with quality score ≥30 and data present in ≥50% of samples. This SNP subset was further filtered to retain only synonymous variants to ensure that estimates reflected putatively neutral diversity. The filtered monomorphic and synonymous SNP VCFs were then merged for downstream analyses.

Nucleotide diversity (π)^61^ was calculated in non-overlapping 5,000 bp windows using the software Pixy^62^, which accounts for missing and invariant sites. Mean π was computed by summing the observed nucleotide differences across all windows and dividing by the total number of comparable (callable) sites. Effective population size (Ne)^63^ was then estimated as: 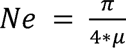 where μ is the mutation rate of 1.7×10^-9^ as defined by Bopp et al., 2013.^64^ To evaluate the influence of sample size on diversity estimates, π was also calculated using a rarefied sample size (n = 18) per time point. Changes in genetic diversity over time were further investigated by calculating minor allele frequencies (MAF) from the synonymous VCF file. For each site, the alternative allele frequency (AF) was compared to the reference allele frequency (1 − AF), and the lesser of the two was recorded as the MAF.

### Simulations

We used two complementary simulation approaches to assess the impact of PAMAFRO and associated reductions in population size on the fixation of *hrp2/3* deletions.

First, we used SLiM^25^ to simulate allele frequency trajectories and changes in parasite population size over a 15-year period, using annual P. falciparum malaria incidence data reported by CDC Peru.^8^ Given the regional heterogeneity of *hrp2/3* deletion prevalence in Peru^42^, we modeled three different populations with varying number of parasites based on ii) total number of cases reported nationwide, ii) total number of cases reported in the province of Maynas, and iii) number of cases reported in the districts most represented in our dataset: Iquitos, Punchana, and San Juan Bautista. These districts account for 790 of 859 samples (92.0%) for which we have *hrp2/3* genotype data. Simulations followed a Wright-Fisher haploid model assuming a 95% cloning rate (i.e., 5% of infections result from outcrossing), consistent with the high level of clonality observed in Peru. We modeled two unlinked loci –*hrp2* and *hrp3*– which reside on separate chromosomes, with free recombination between them. The average frequencies of four genotypes observed in 2003–2004 were used to initialize the ancestral population: *hrp2-/3-*, *hrp2-*, *hrp3-*, and *hrp2+/3+* (no deletions). No back mutations or new mutations were introduced during simulations, which is reasonable given the short evolutionary timescale. Demographic trajectories were modeled over 135 generations (assuming 9 *P. falciparum* generations per year, based on an estimated two-month generation time^65^) from 2003 to 2018. Final allele frequencies were extracted at generation 135 (year 2018), and each simulation was repeated 1,000 times per scenario. Selection coefficients were assumed to be equal for the two deletions and were applied multiplicatively, such that the relative fitnesses were *w_WT_* = 1, *w_hrp_*_2-_ = (1 + *s*), *w_hrp_*_3-_ = (1 + *s*), and *w_hrp_*_2-/3-_ = (1 + *s*)^2^. Simulations were run across selection coefficients (*s* = 0.0, 0.001, 0.01, 0.03, 0.05, 0.07,0.1). For each region and selection coefficient, 1000 replicate simulations were performed.

In addition, we used a previously published *P. falciparum* transmission modelling framework that accounts for age-related immunity, heterogeneous mosquito biting rates, treatment, and parasite dynamics, and which has been used extensively to model the emergence and spread of hrp2/3 deletions.^14,26^ Building on the same underlying transmission model used in these earlier HRP2 studies, we used the implementation in the *magenta* individual-based model.^66^ This model extends the previously used models by modelling both individual mosquitoes and parasite clones (depicted as barcodes encoding for different genotypes at different genome positions), explicitly capturing the full parasite life cycle and enabling recombination and the generation of new parasite clones through recombination to be tracked. Using *magenta,* we first calibrated malaria transmission intensity to match pre-PAMAFRO per-capita *P. falciparum* case incidence in the three districts most represented in our dataset (Iquitos, Punchana, and San Juan Bautista).^8^ We then ran 500 forward-time simulations across a range of human population sizes (25,000–200,000) for 30 years to ensure steady-state equilibrium. Following this burn-in period, parasite haplotype frequencies were reinitialised such that population-level *hrp2/3* frequencies matched the average observed between 2003–2006. To model the impact of PAMAFRO, we simulated a sustained reduction in malaria transmission for 4 years followed by a sustained increase in transmission for 8 years, again calibrated to the per-capita malaria incidence observed during and after PAMAFRO. From each simulation, we calculated the proportion of runs in which *hrp2-/3-* reached an average frequency greater than 75% during 2016–2018, which we defined as fixation, and estimated Binomial confidence intervals for each proportion. To directly estimate the effect of PAMAFRO on deletion frequency, we also ran parallel simulations without the PAMAFRO intervention, maintaining constant transmission intensity throughout. We fit a Binomial regression model to the simulation outputs, modelling fixation as a function of population size and its interaction with the presence of PAMAFRO. From the model, we reported odds-ratios of the impact of PAMAFRO on changing population size and the absolute fixation probabilities resulting from PAMAFRO at various population sizes reflective of the population size of the three districts as estimated by WorldPop for 2005 (i.e. prior to PAMAFRO).^67,68^

## Supporting information

Supplemental results

## Data and code availability

All MIP sequencing data generated and analyzed in this study have been deposited in NCBI SRA under project number PRJNA1370144. Custom scripts used for data processing and analysis are available on GitHub at https://github.com/isabelagyuricza/Peru_Pfalciparum_PAMAFRO. SLiM simulation scripts are available at https://github.com/JohriLab/PfalciparumPeruSims. Simulations scripts for the malaria transmission model are available at https://github.com/OJWatson/peru_hrp23.^69^

## ACKNOWLEDGEMENTS

We are grateful to the research teams at Universidad Peruana Cayetano Heredia, NAMRU SOUTH, and the GERESA Loreto for their efforts in conducting fieldwork, and to all the participants and their family members who generously contributed to this study. We thank Ruthly François-Zafka, Rebecca Crudale, and Alec Leonetti for training and guidance in performing MIP experiments. We also thank Oksana Kharabora for supporting the laboratory work. This work was supported by the National Institute of Allergy and Infectious Diseases (R01AI177791 to JBP and K24AI134990 to JJJ), the National Institute of General Medical Sciences of the National Institutes of Health (R35GM154969 to PJ), the Global Emerging Infections Surveillance (GEIS) Branch (PROMIS ID P0146_25_N6 to HOV) of the Defense Health Agency’s Armed Forces Health Surveillance Division (AFHSD).

## DISCLAIMER

The views expressed in this article reflect the results of research conducted by the authors and do not necessarily reflect the official policy or position of the Department of the Navy, Department of Defense, nor the U.S. Government.

## COPYRIGHT STATEMENT

Some authors of this manuscript are military service members or employees of the U.S. Government. This work was prepared as part of their official duties. Title 17 U.S.C. §105 provides that Copyright protection under this Title is not available for any work of the United States Government. Title 17 U.S.C. §101 defines a U.S. Government work as a work prepared by a military service member or employee of the U.S. Government as part of that person’s official duties.

## ETHICS APPROVAL AND CONSENT TO PARTICIPATE

The study protocol NMRCD.2007.0004 was approved by the U.S. Naval Medical Research Unit SOUTH Institutional Review Board in compliance with all applicable Federal regulations governing the protection of human subjects.

## DECLARATION OF GENERATIVE AI AND AI-ASSISTED TECHNOLOGIES IN THE MANUSCRIPT PREPARATION PROCESS

During the preparation of this work, the authors used ChatGPT to assist with text editing and to identify and address coding errors. After using this tool/service, the authors reviewed and edited the content as needed and take full responsibility for the content of the published article.

