## Supplemental results for "Malaria control and the unexpected spread of diagnostic-resistant *Plasmodium falciparum* in Peru"

**SUPPLEMENTAL TEXT**

We genotyped samples for *hrp2/3* deletion status using two approaches: multiplexed real-time PCR, which detects *hrp2/3* via amplification of exon 2, and conventional PCR. Multiplexed real-time PCR was performed in our laboratory (**Supplemental File 1**) and adapted from Grignard et al.^1^ Controls included *P. falciparum* lactate dehydrogenase (*pfldh*) and human beta-tubulin (*Hbb*) genes, along with laboratory strains 3D7, DD2, and HB3 as standards, as previously described.^1^ To evaluate assay reproducibility, one qPCR plate was run in duplicate, resulting in a 16% (47 of 288 samples) discordance rate between the two runs. Most discordant calls (75%; 35 of 47 samples) were due to indeterminate results arising from either *pfldh*/*Hbb* amplification failure or inadequate Ct values (Ct < 15 or Ct > 35) for deletion calling (**Supplemental Table 2**). *Hrp2/3* deletion calls were made for 65% (790 out of 1,215) of samples (**Supplemental Table 3**). Amplification results for controls are shown in **Supplemental Table 4**. Additionally, previously published conventional PCR results from collaborators allowed us to assign *hrp2/3* deletion status to an additional 69 samples, achieving a total of 859 samples (70.7%) with final deletion calls (**Supplemental Table 1**). From the 320 samples with both real-time and conventional PCR data, 51 (15.9%) had a discordant *hrp2/3* deletion call between the two methods. For those, we considered the real-time PCR result as the final deletion call. We also observed high concordance between *hrp2/3* genotyping by real-time PCR and MIP sequencing. Of the 123 samples with sufficient coverage for chromosomes 8 and 13, only two (MDP4208 and MDP6027) were misclassified by PCR as having *hrp3* deletions only (*hrp3-*), despite lacking reads for both *hrp2* and *hrp3* in the MIP data.

**SUPPLEMENTAL FIGURES**

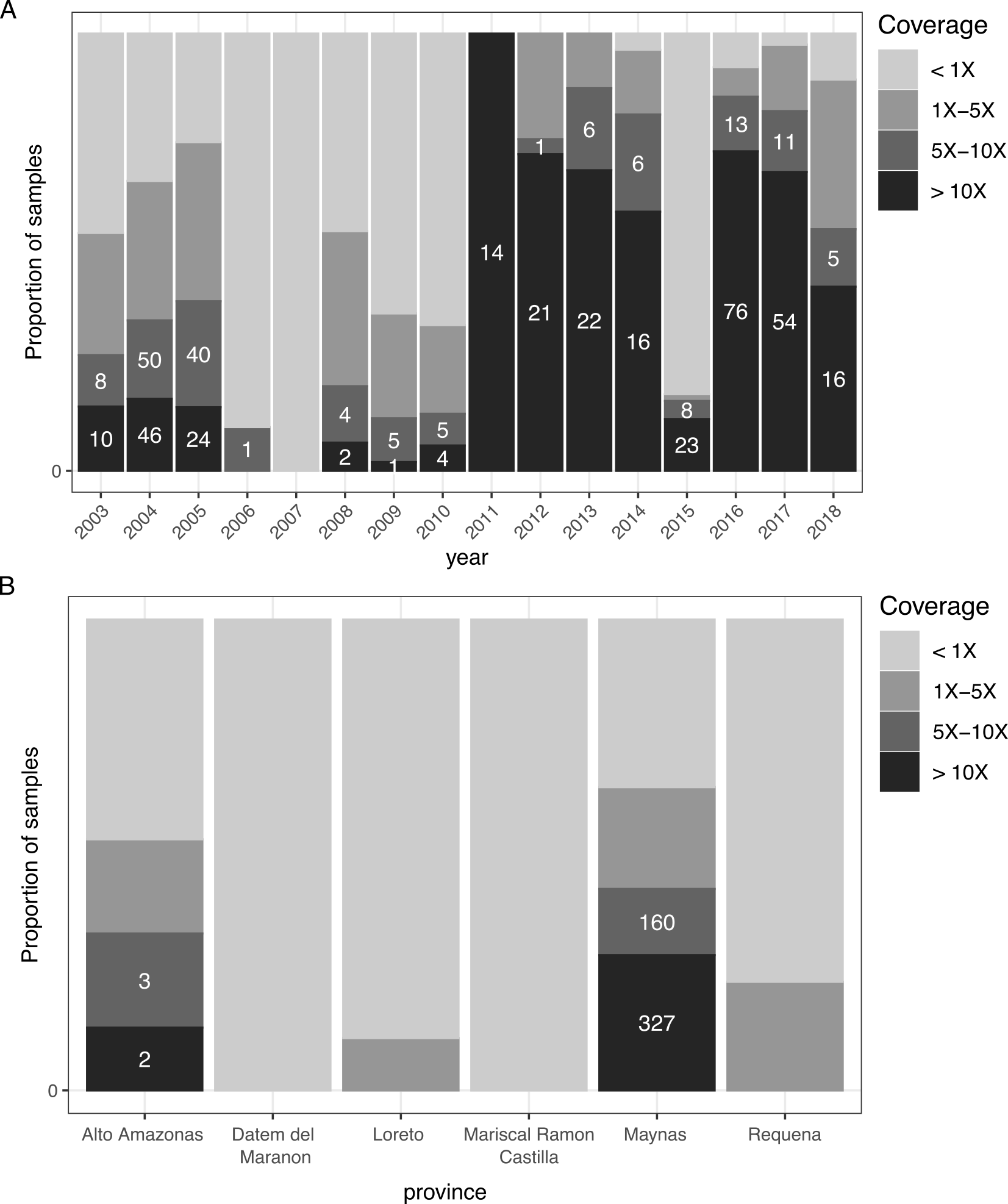

**Supplemental Figure 1. Genome-wide MIP sequencing coverage by year.** Coverage for each *P. falciparum* sample (n = 1,215) was estimated by dividing the total number of unique molecular identifiers (UMI) counts by the total number of MIP probes in the panel (n = 2,128). The bars represent the proportion of parasites of each coverage category by (A) year and (B) province. Only samples with mean coverage ≥5X (average of ≥5 UMIs / probe) were retained for downstream analysis. The number of samples meeting this threshold per year is indicated within each bar.

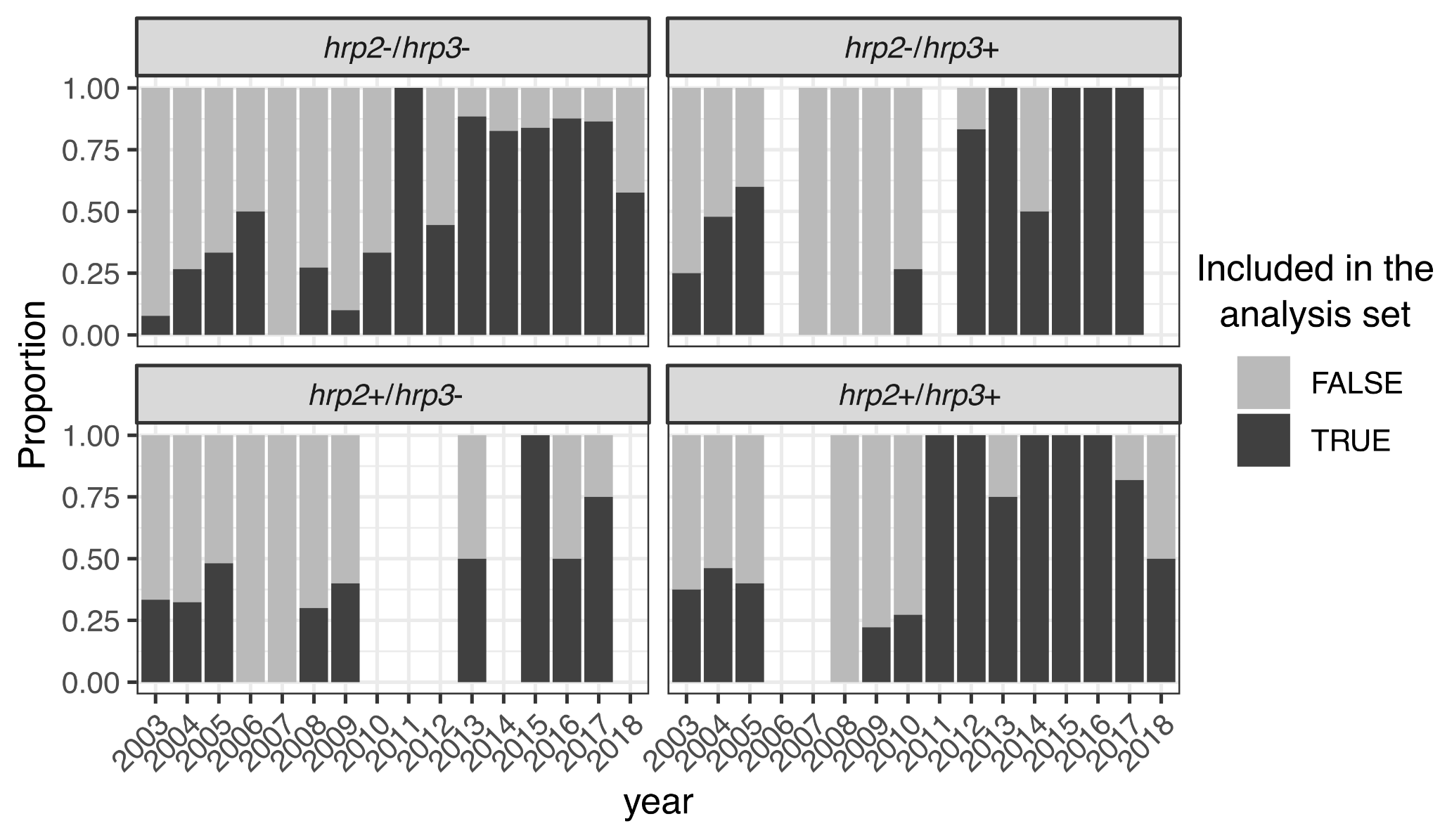

**Supplemental Figure 2. Genome-wide MIP sequencing capture success by *hrp2/3* genotype and year**. Proportion of *P. falciparum* samples (n = 1,215) retained for downstream analysis, defined as those meeting the ≥5X genome-wide MIP coverage threshold. Samples are grouped by year and faceted by *hrp2/3* genotype. Bars represent the proportion of successfully captured samples (black) versus those that did not meet the threshold (gray), highlighting genotype-specific and temporal gaps in sequencing coverage.

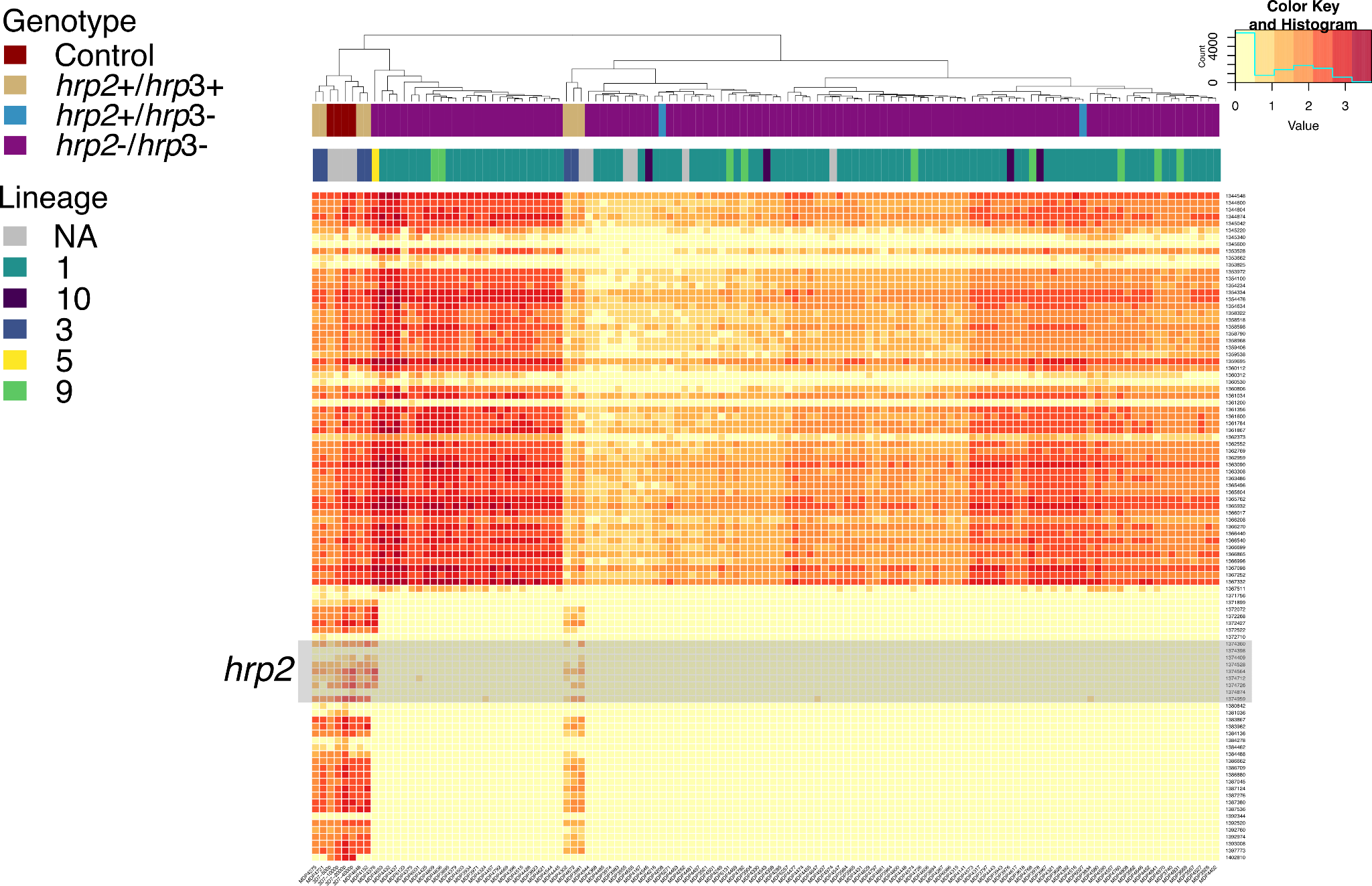

**Supplemental Figure 3. *hrp2/3*-targeted MIP sequencing coverage across chromosome 8.** Heatmap of unique molecular identifier (UMI) read counts for chromosome 8, including the *hrp2* gene (highlighted by a gray box) and its flanking regions for n = 123 samples with high-quality data. Read counts are log₁₀-transformed. Chromosomal positions (y-axis) are ordered from centromere to telomere, and parasite samples (x-axis) are sorted by hierarchical clustering determined by the R function *hclust*(). The top annotation bar indicates *hrp2/hrp3* genotype category; no *hrp2–/hrp3+* samples were successfully captured in the *hrp2/3*-targeted MIP panel. The bottom annotation bar denotes parasite lineage, as defined by identity-by-descent (IBD) analysis using a separate MIP panel with targets across the genome. Samples labeled as “NA” were excluded from the IBD analysis due to insufficient genome-wide MIP coverage.

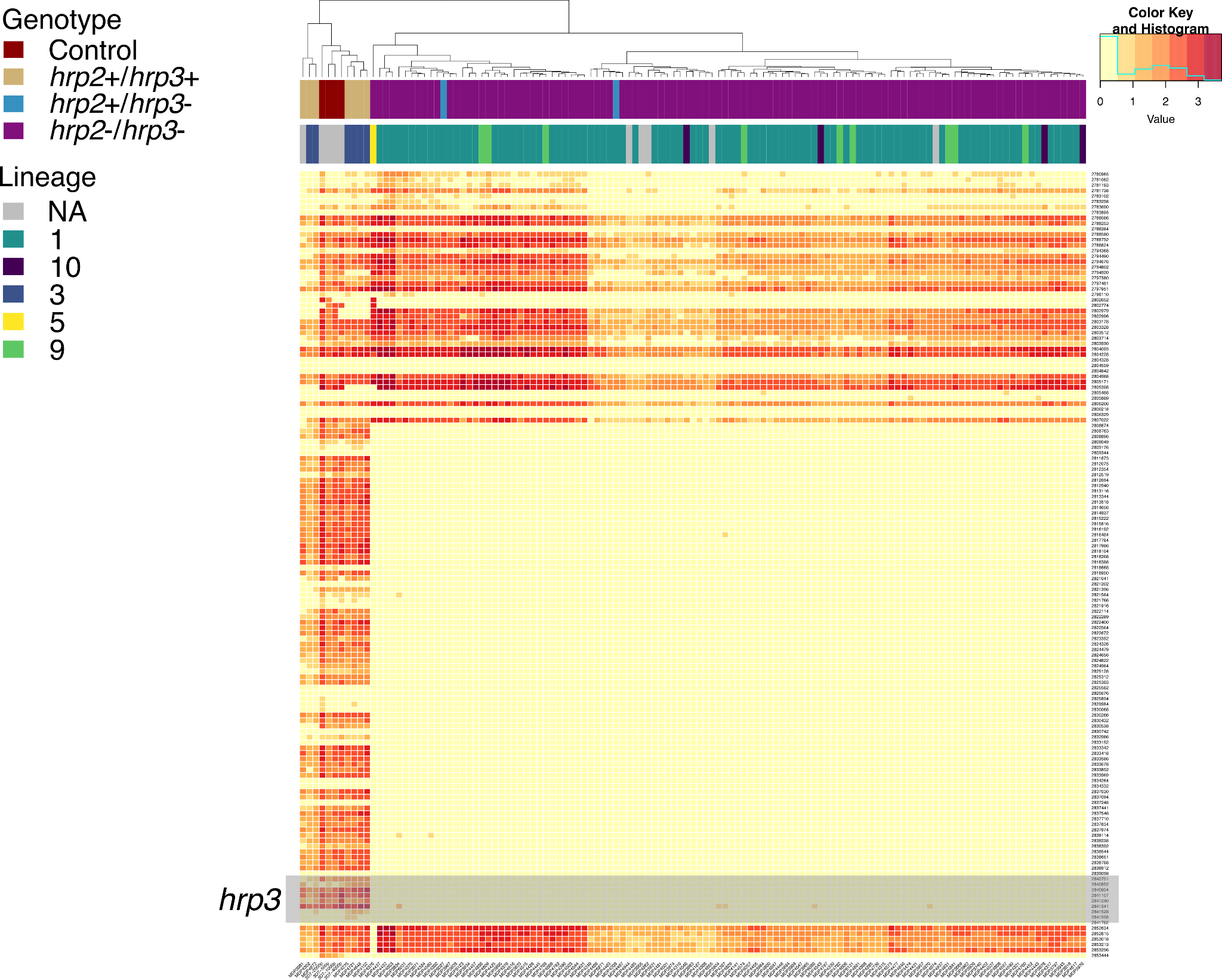

**Supplemental Figure 4. *hrp2/3*-targeted MIP sequencing coverage across chromosome 13.** Heatmap of unique molecular identifier (UMI) read counts for chromosome 13, including the *hrp3* gene (highlighted by a gray box) and its flanking regions for n = 123 samples with high-quality data. Read counts are log₁₀-transformed. Chromosomal positions (y-axis) are ordered from centromere to telomere, and parasite samples (x-axis) are sorted by hierarchical clustering determined by the R function *hclust*(). The top annotation bar indicates *hrp2/hrp3* genotype category; no *hrp2–/hrp3+* samples were successfully captured in the *hrp2/3*-targeted MIP panel. The bottom annotation bar denotes parasite lineage, as defined by identity-by-descent (IBD) analysis. Samples labeled as “NA” were excluded from the IBD analysis due to insufficient genome-wide MIP coverage. The reads aligned observed at the distal end of the chromosome represent multimapping reads arising from a segmental duplication shared between chromosomes 13 and 11, as previously described by Hathaway et al.^2^

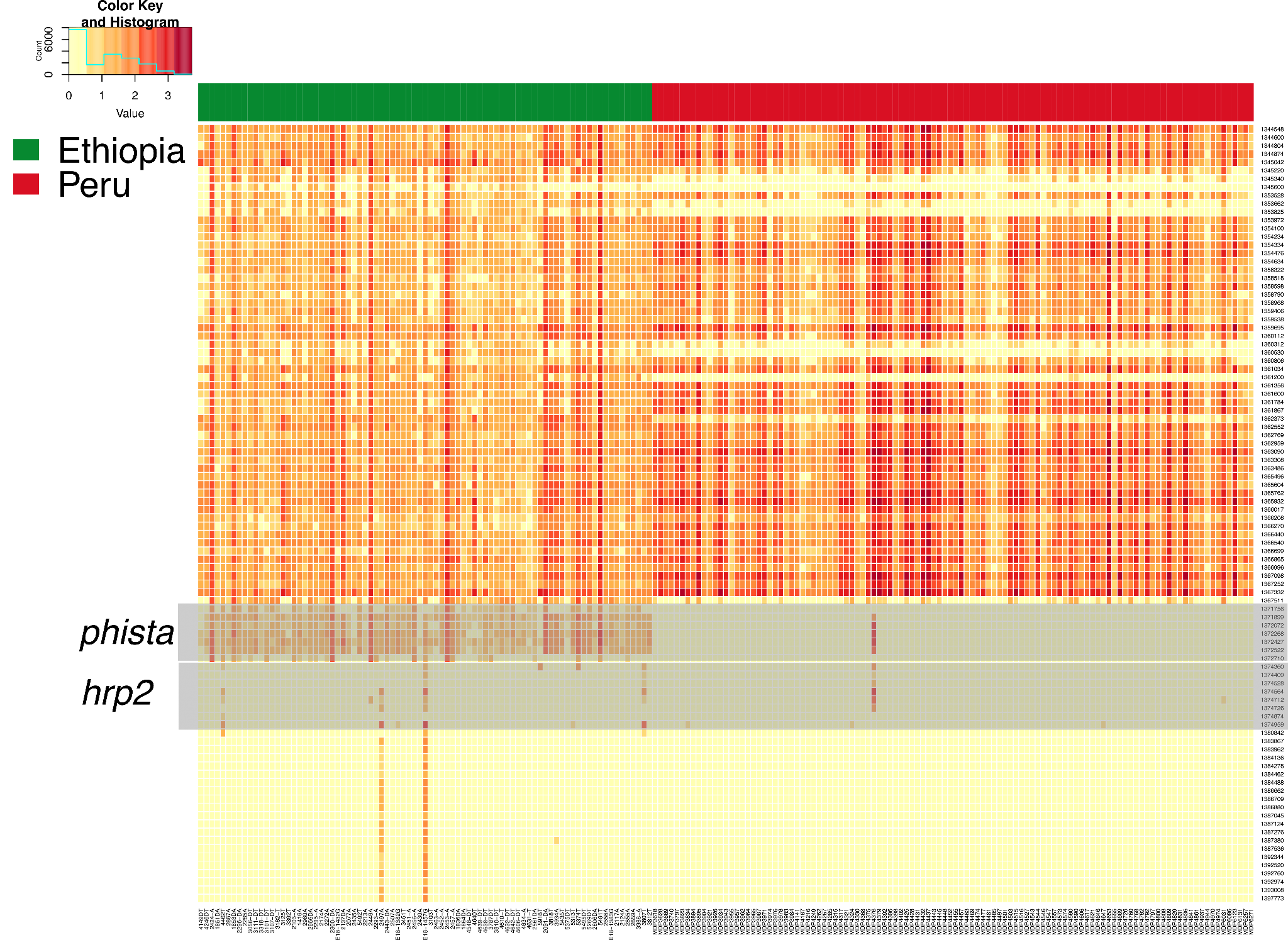

**Supplemental Figure 5. *hrp2/3*-targeted MIP sequencing coverage across chromosome 8 between Peru and Ethiopia.** Heatmap of unique molecular identifier (UMI) read counts for chromosome 8 for n = 167 (Ethiopia) and n = 123 (Peru) high-quality (coverage ≥5X) samples. Read counts are log₁₀-transformed. Chromosomal positions (y-axis) are ordered from centromere to telomere, and parasite samples (x-axis) are grouped by country of origin. Only samples annotated as *hrp2-/hrp3+* or *hrp2-/hrp3-* according to multiplexed PCR are shown. Gray boxes highlight the locations of the *hrp2* and *phista* genes. Samples that show coverage for *hrp2* are likely incorrectly classified in the PCR. Notably, coverage is observed over *phista* in Ethiopian parasites but absent in Peruvian samples, illustrating that the *hrp2/3*-targeted MIP panel can resolve deletion breakpoint differences between these populations.

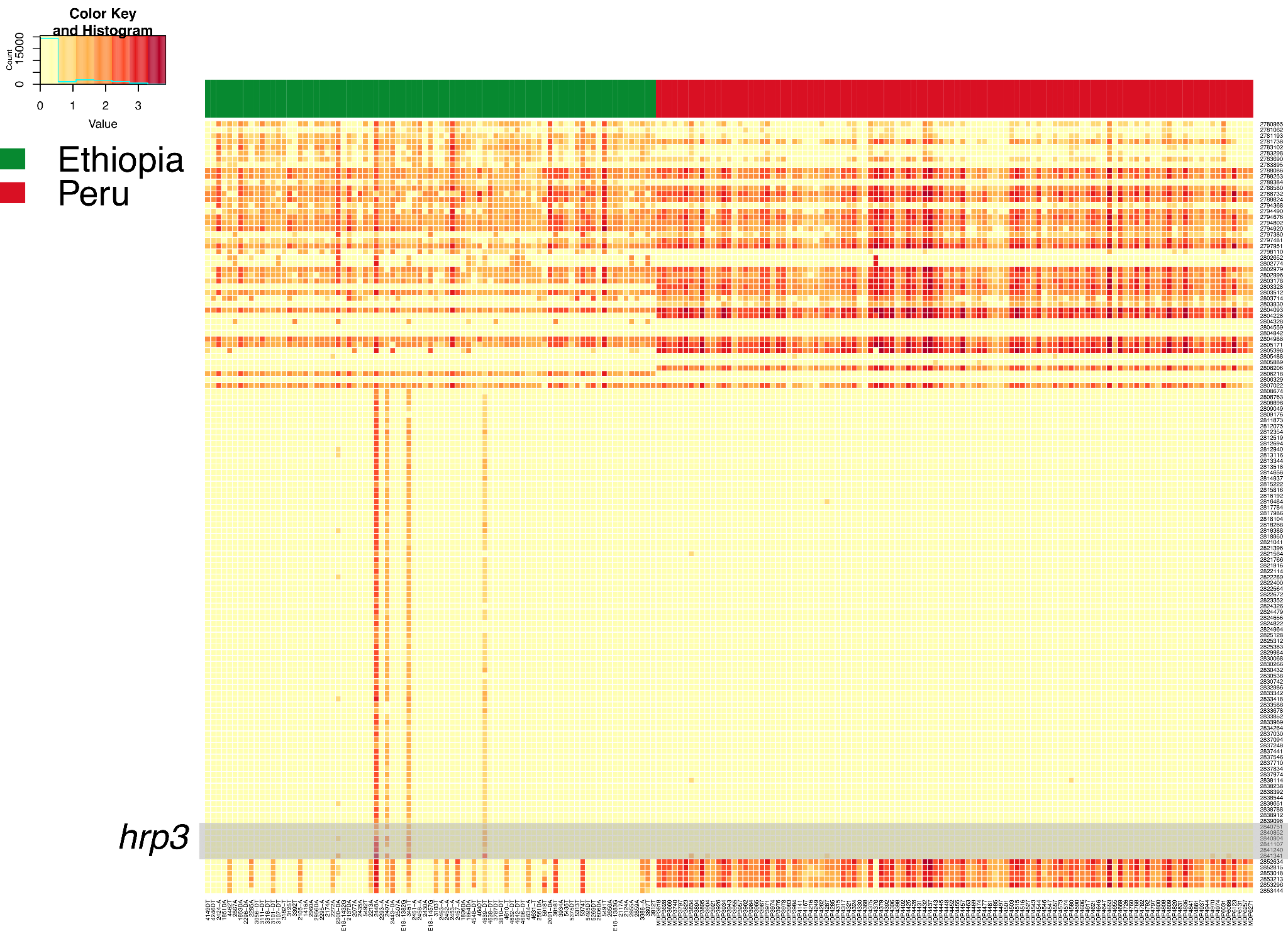

**Supplemental Figure 6. *hrp2/3*-targeted MIP sequencing coverage across chromosome 13 between Peru and Ethiopia.** Heatmap of unique molecular identifier (UMI) read counts for chromosome 8 for n = 167 (Ethiopia) and n = 123 (Peru) high-quality (coverage ≥5X) samples. Read counts are log₁₀-transformed. Chromosomal positions (y-axis) are ordered from centromere to telomere, and parasite samples (x-axis) are grouped by country of origin. Only *hrp2+/hrp3*- or *hrp2-/hrp3-* parasites are shown. Gray boxes highlight the locations of the *hrp3* gene. Chromosomal breakpoints between Ethiopian and Peruvian parasites are similar, and the reads at the distal end of the chromosome represent multimapping reads arising from a segmental duplication shared between chromosomes 13 and 11, as previously described by Hathaway et al.^2^

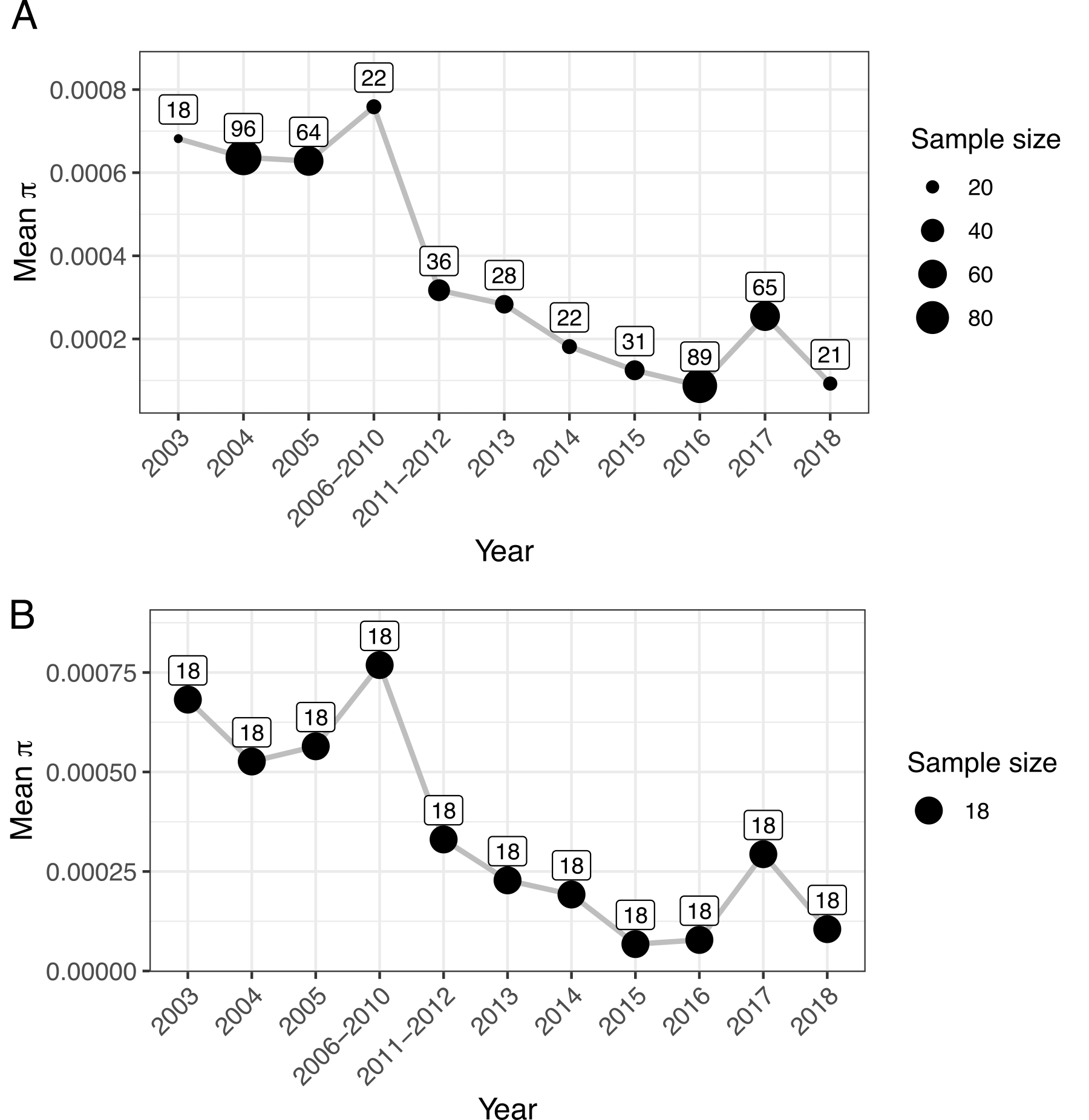

**Supplemental** **Figure 7. Nucleotide diversity (π)** **over time**. Mean π was calculated across non-overlapping 5 kb windows using synonymous and invariant sites (n = 492). For each year group, π was computed by summing observed nucleotide differences across all windows and dividing by the total number of callable sites. a) Mean π per year, based on all available samples meeting quality thresholds. Years with fewer than 18 samples were combined with adjacent years to ensure sufficient representation. Point size reflects sample size, and exact counts are shown. b) Mean π per year using a rarefied dataset of 18 randomly selected samples per year to assess the effect of varying sample sizes on π estimation.

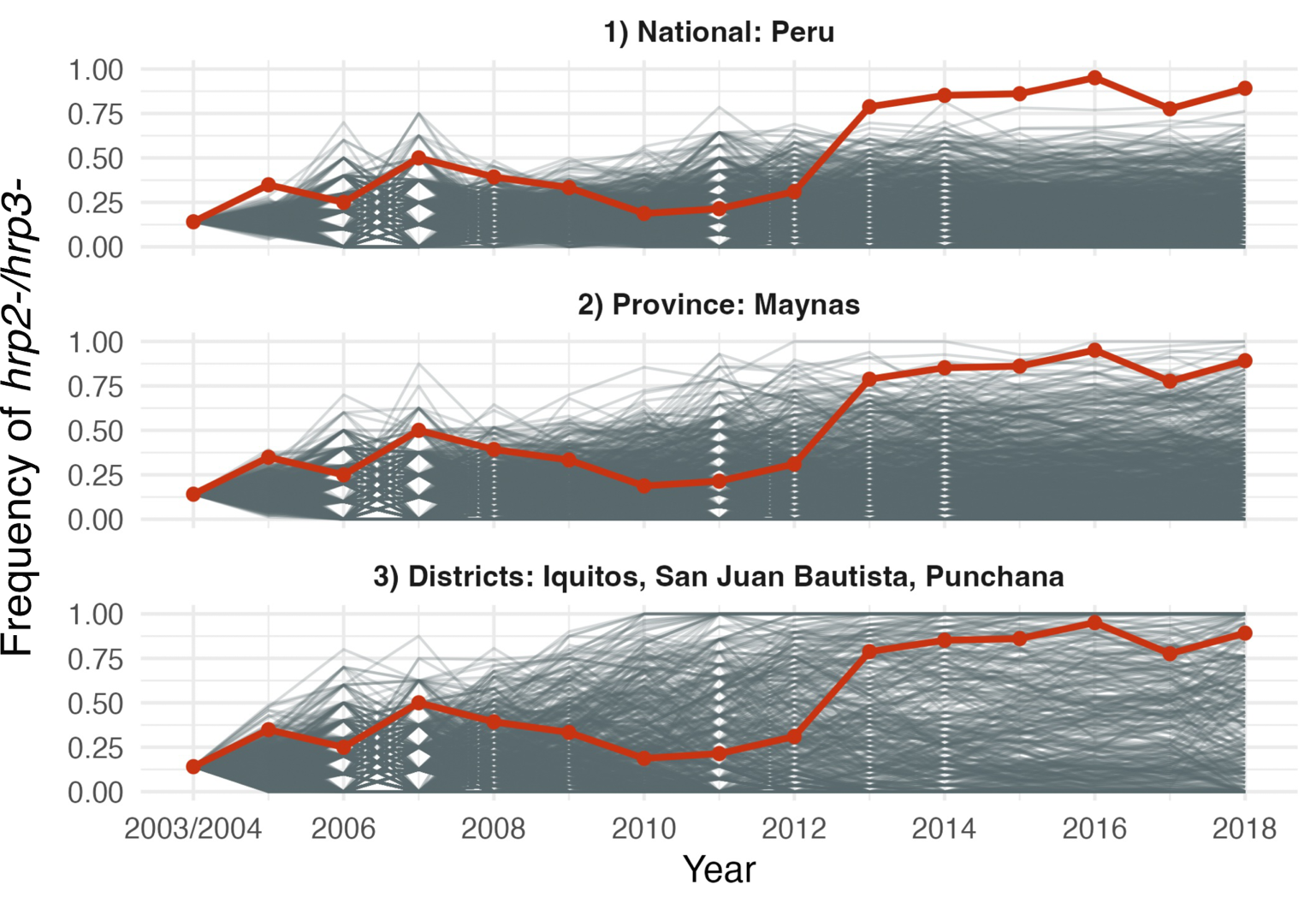

**Supplemental** **Figure 8. SLiM allele trajectories of *hrp2-/3-* over time.** Simulated (grey) *versus* observed (red) frequencies of *hrp2-/3-* over time. A total of 1,000 simulations were conducted under neutrality (s = 0) based on annual *P. falciparum* case reports from CDC Peru for three spatial scales: (i) all reported cases nationwide (National); (ii) reported cases for the province of Maynas (Province); and (iii) reported cases from Iquitos, Punchana, and San Juan Bautista (Districts). Differences between observed and simulated *hrp2-/3-* frequencies (assuming *s* = 0) become prominent after 2012.

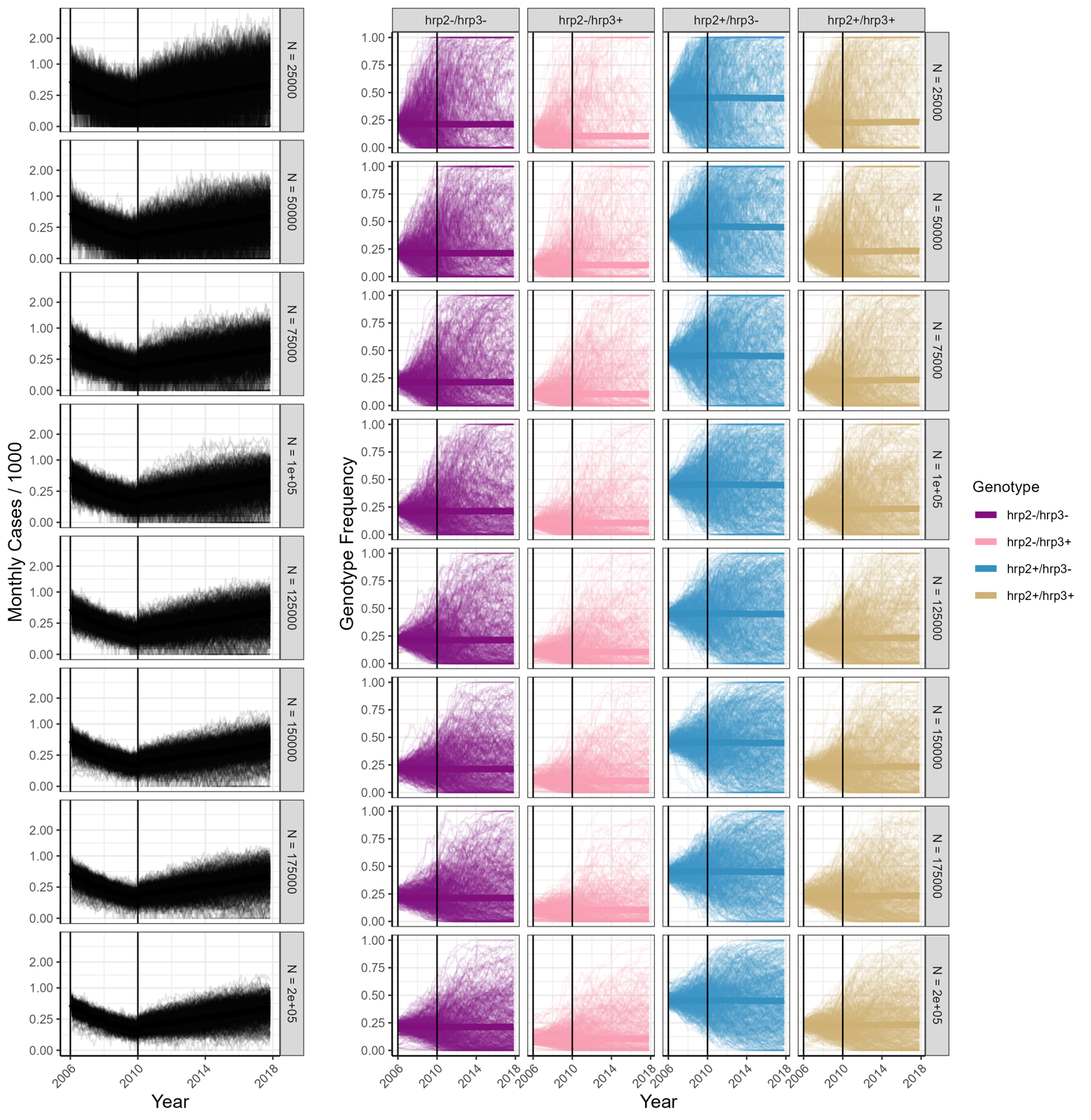

**Supplemental Figure 9.** **Allele trajectories of *hrp2/3* deletions over time according to *P. falciparum* malaria transmission model.** Monthly number of malaria cases per 1000 individuals over time for different human populations varying from 25,000 to 200,000 individuals. A total of 500 simulations (thinner lines) were conducted, and the thicker lines represent the median across simulations. Smaller human populations show the highest stochasticity in allele frequencies and, consequently, probability of increased *hrp2-/3-* frequency over time due to drift.
